# MCM Double Hexamer Loading Visualised with Human Proteins

**DOI:** 10.1101/2024.04.10.588848

**Authors:** Florian Weissmann, Julia F. Greiwe, Thomas Pühringer, Thomas C. R. Miller, John F. X. Diffley, Alessandro Costa

**Affiliations:** Chromosome Replication Laboratory, The Francis Crick Institute, London NW1 1AT, U.K; Macromolecular Machines Laboratory, The Francis Crick Institute, London NW1 1AT, U.K; DNRF Center for Chromosome Stability, Department of Cellular and Molecular Medicine, University of Copenhagen, Copenhagen, Denmark

## Abstract

Eukaryotic DNA replication begins with the loading of the MCM replicative DNA helicase as a head-to-head double hexamer (DH) at origins of DNA replication^1–3^. Our current understanding of how DH is assembled by the Origin Recognition Complex (ORC), CDC6 and CDT1 comes mostly from budding yeast. Here we characterise human DH (hDH) loading using biochemical reconstitution and cryo-electron microscopy with purified proteins. We show that hDH engages DNA differently from yeast (yDH), and generates ∼5 base pairs of unwound DNA at the interface between hexamers, as seen in hDH isolated from cells^4^. We identify several differences from yeast in the order of factor recruitment and dependencies during hDH assembly. Unlike yeast^5–8^, the ORC6 subunit of ORC is not essential for initial MCM recruitment or hDH loading, but contributes to an alternative hDH assembly pathway requiring an intrinsically disordered region (IDR) in ORC1, which may work through a novel MCM-ORC (hMO*) intermediate. Our work presents a detailed view of how DHs are assembled in an organism utilising sequence-independent replication origins, it provides further evidence for diversity in eukaryotic DH assembly mechanisms^9^, and it represents the first step toward reconstitution of DNA replication initiation with purified human proteins.

## Results

### Reconstitution of MCM Loading with Purified Human Proteins

We expressed human ORC, CDC6, CDT1 and MCM using the biGBac baculovirus expression system^10^ (Fig. 1a). Consistent with previous work^11,12^, we found that ORC6 did not co-purify with ORC1-5 when co-expressed, so we produced ORC1-5 and ORC6 separately. Similarly, unlike yeast, CDT1 did not co-purify with MCM, so CDT1 and MCM were expressed and purified separately. We previously developed approaches to examine yDH loading using DNA coupled to magnetic beads, which allowed us to isolate yDH and examine its properties, for example, its stability after high salt wash^7,8,13,14^. With this approach and the human proteins, we were able to detect ATP- and DNA-dependent recruitment of ORC and MCM to bead-bound DNA, but complexes were not resistant to high salt extraction and did not require components including ORC6 and CDC6 (ED Fig. 1a,b).

**Figure 1.**
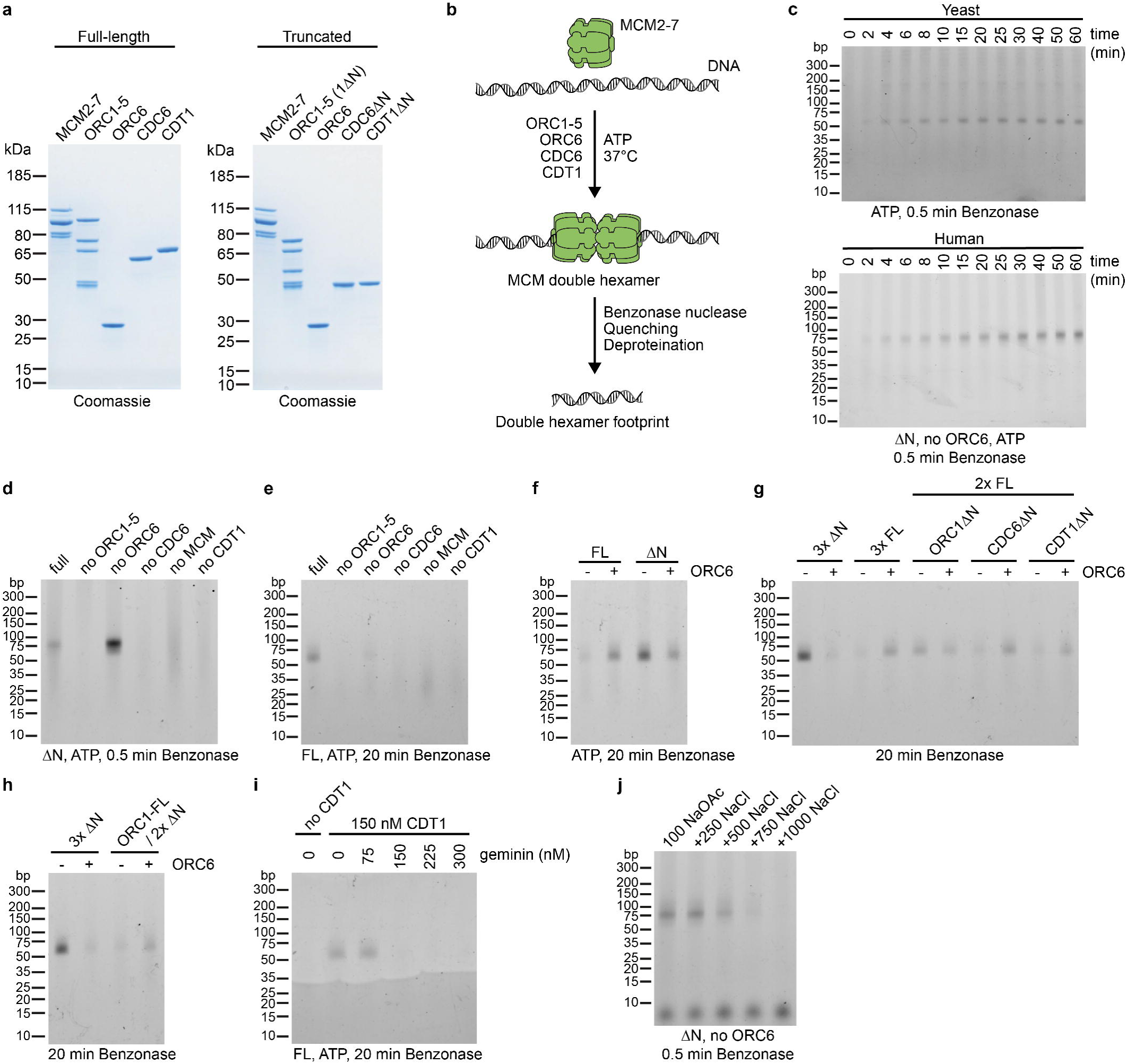
Reconstitution of human DH loading. **a**, Purified human MCM loading proteins analysed by SDS-PAGE and Coomassie staining. Full-length proteins (left, FL) and truncated proteins (right, ΔN): ORC1ΔN, CDC6ΔN, CDT1ΔN. **b**, Outline of the nuclease footprinting assay. MCM loading reactions are treated with Benzonase followed by quenching with EDTA, SDS and proteinase K. Then DNA is purified by phenol-chloroform-isoamyl alcohol extraction and ethanol precipitation, resolved on a TBE polyacrylamide gel, and stained with SYBR Gold. **c**, Timecourse of yeast (top) and human (bottom) MCM loading reactions. **d**, Protein requirements for the DH footprint with truncated human proteins. **e**, Requirements of MCM loading with full-length proteins. **f**, Side-by-side comparison of the ORC6 dependency with full-length and truncated proteins. **g**, As f, testing conditions in which either ORC1, CDC6, or CDT1 was truncated and other proteins full-length. **h**, As f, with full-length ORC1 and truncated CDC6 and CDT1. **i**, Effect of geminin on full-length MCM loading reactions. CDT1 and geminin were pre-mixed before reactions were started. **j**, Salt stability of the DH. MCM was loaded for 30 minutes and then incubated in buffers containing the indicated concentrations of sodium chloride for 15 minutes, followed by dilution and Benzonase treatment.

Nonetheless, negative-stain EM imaging of a human MCM loading reaction on a short synthetic yeast origin flanked by nucleosomes yielded 2D averages similar to yeast DH (ED Fig. 1c), suggesting that hDH was being assembled, but that we might need more informative biochemical assays. The N-terminal IDRs of the *Drosophila melanogaster* orthologues of ORC1, CDC6 and CDT1 drive liquid-liquid phase separation^15,16^, which might contribute to high background in bead-based assays and interfere with DH assembly *in vitro* with human proteins as well, so we wanted a biochemical assay that does not require bead-bound DNA.

We therefore developed a nuclease footprinting assay (Fig. 1b) to examine DNA-bound protein complexes. In this assay, complexes are assembled on soluble plasmid DNA and DNA is digested extensively with Benzonase; all that remains after digestion is the DNA protected by proteins, which is resolved on polyacrylamide gels. With yeast ORC, CDC6 and MCM-CDT1 in the presence of ATP, a band of ∼60 base pairs (bp) — approximately the length of DNA contained within the central channel of the yeast DH^17^ — appeared with time, beginning at 2-4 min (Fig. 1c, top). We expressed and purified human ORC, CDC6 and CDT1 lacking the N-terminal IDRs of ORC1, CDC6 and CDT1(‘truncated proteins’, Fig. 1a) to avoid complications from liquid-liquid phase separation. Using these truncated proteins in the footprinting assay we saw a band of slightly larger size (∼75bp) that appeared with similar kinetics (Fig. 1c, bottom). This assay therefore detects the hDH very similar to the yDH.

The appearance of the hDH required ORC1-5, CDC6, MCM and CDT1, but did not require ORC6 according to DNA footprinting (Fig. 1d) and negative stain EM (ED Fig. 2); in fact, the presence of ORC6 consistently reduced its amount. With the full length proteins, we saw a band of the same size that also required ORC1-5, CDC6 and CDT1; in this case, however, only a faint band was seen when ORC6 was omitted (Fig. 1e). Fig. 1f shows MCM loading by the truncated and full length proteins side-by-side, emphasising that ORC6 stimulates DH assembly with full-length proteins, but inhibits assembly with the truncated proteins. From this we conclude that there is a pathway for MCM loading that does not require ORC6 and that this is the preferred pathway when truncated proteins are used. The ORC6 stimulation with full length proteins suggests there is an alternative ORC6-dependent MCM loading pathway that requires one or more of the IDRs. To determine which IDR is responsible for this ORC6 effect, we tested each truncation alone. Fig. 1g shows that ORC6 stimulated loading with either CDC6ΔN or CDT1ΔN; however with ORC1ΔN, ORC6 no longer stimulated loading but instead inhibited it. The inclusion of full length ORC1 with CDC6ΔN and CDT1ΔN conferred ORC6-dependence (Fig. 1h), indicating that the ORC1 IDR is necessary and sufficient for the ORC6 dependent pathway.

To characterise MCM loading further, we next tested the effect of geminin, which inhibits hDH assembly *in vivo* by inhibiting CDT1^18–20^. Consistent with this, MCM loading *in vitro* by both full length and truncated proteins was inhibited by geminin, and inhibition occurred when geminin levels were equal to or greater than the level of CDT1 (Fig. 1i). To examine salt stability of hDH, reactions were challenged with different salt concentrations, then reactions were diluted and benzonase treated. In these reactions, hDH was stable up to 500 mM NaCl, but not above. In fact, hDH was stable in 500 mM NaCl (Fig. 1j) for up to 60 min (ED Fig. 1d); hDH could also be detected after 750 mM NaCl treatment for short time periods, but diminished in a time dependent manner (ED Fig. 1e) indicating the human DH is less salt-stable than the yeast DH, which is stable up to 2 M NaCl^21^. Human ORC is not a sequence specific DNA binding protein^22–24^, and consistent with this, MCM loading *in vitro* was similarly efficient on DNA containing or lacking human replication origin sequences (ED Fig. 1f).

### The Human MCM Double Hexamer

To characterise hDH and to understand its interactions with DNA, we used cryo-EM to image the human MCM loading reaction earlier visualised by negative-stain EM. We solved a 3.1 Å resolution structure of the hDH (ED Fig. 3). The two MCM hexamers, related by two-fold symmetry, interact via two N-terminal tiers forming a tilted homodimerisation interface that constricts the MCM central channel. The inter-ring register matches that observed in the yDH structure^25^, with the two MCM2-5 subunits mapping on opposite sides in the hDH (Fig. 2a). The active site at the interface between MCM6 and 2 is ATP bound, the MCM4-7 site is empty and all other sites are ADP bound (Fig. 2b). Our structure of the reconstituted hDH matches that of hDH isolated from cells, although four of the six catalytic sites were bound by ATP or a mixture of ATP and ADP in the structure of hDH from cells^4^. This discrepancy might reflect differences in sample preparation. While in our reconstitution experiment cryo-EM grids were flash-frozen within 45 min of establishing the reaction, hDHs from cells were subjected to purification steps in ATP-containing buffer, which might have favoured nucleotide exchange. Differences in nucleotide occupancy between the two structures, however, show no effect on the protein-DNA contacts.

**Figure 2.**
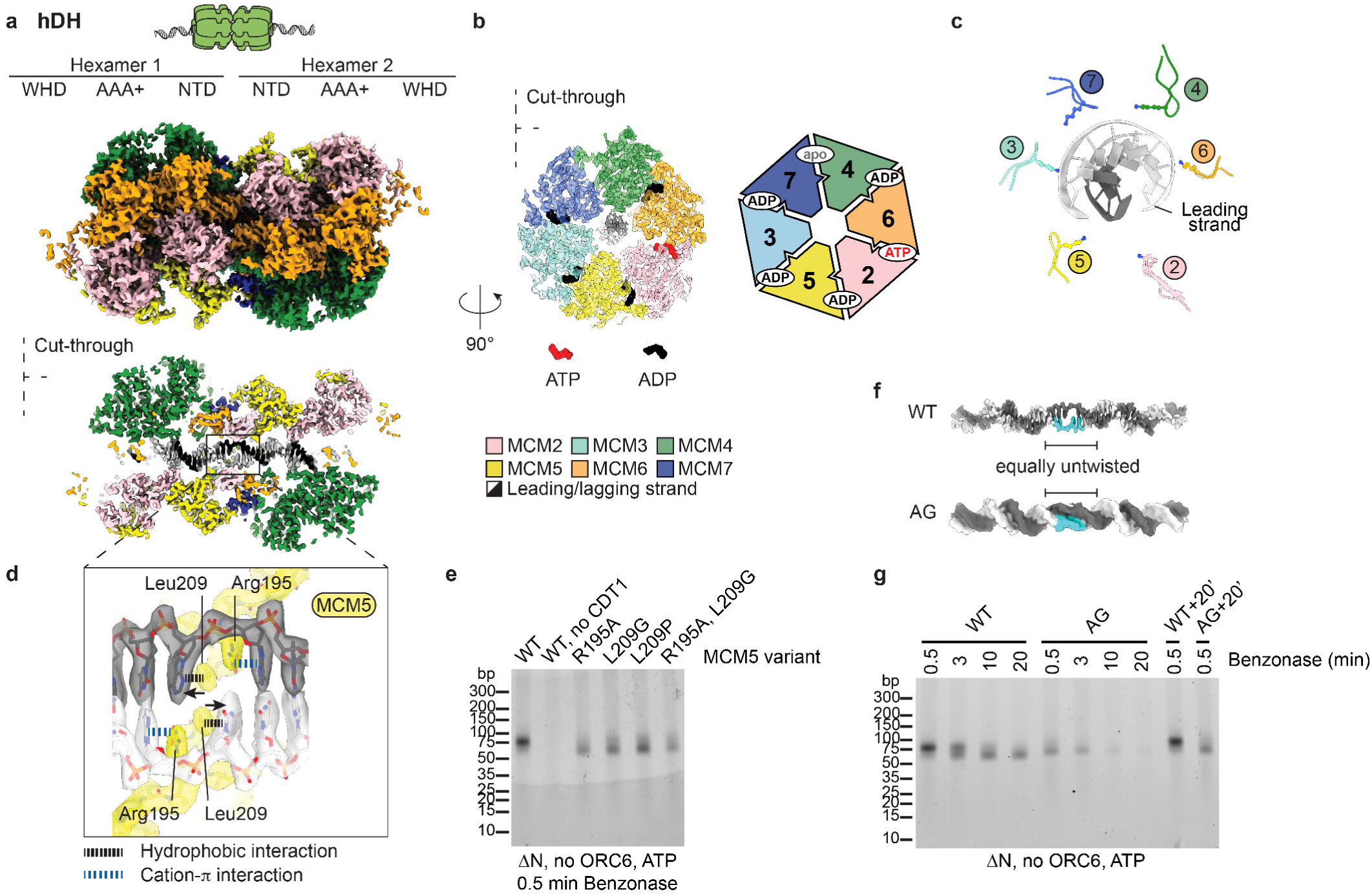
Human DH loading untwists and melts DNA. **a**, Surface rendering and cut-through view of the human MCM double hexamer loaded onto duplex DNA. The double helix is untwisted and one base pair is broken at the dimerisation interface. This distortion allows DNA to traverse a narrow passage created by two MCM5 ZnF domains. **b**, C-terminal ATPase view of one MCM ring. Nucleotides are shown in the ATPase sites between MCM subunits. **c**, PS1 pore loops form a staircase arrangement that follows the leading strand template. **d**, Zoomed-in view of the broken base pair stabilised by R195 and L209 of Mcm5. **e**, Nuclease footprinting assay with MCM5 mutants. **f**, Comparison between duplex DNA in the wild type and MCM5 AG mutant double hexamer reveals a similar degree of DNA untwisting. **g**, MCM WT and AG mutant (MCM5 R195A, L209G) were loaded on DNA and subjected to a timecourse of Benzonase treatment between 0.5 and 20 minutes. Controls were incubated for an additional 20 minutes followed by 0.5 min Benzonase treatment.

45 bp could be clearly resolved inside the MCM hexamer, though the length of the central channel is compatible with a 75 bp protection seen by DNA footprinting. The pre-sensor 1 (PS1) pore loops, emanating from the ATPase modules, are arranged in a staircase configuration that follows the DNA double helix. PS1 pore loops of MCM7,4,6 engage in minor groove contacts with the strand running 3’ to 5’ from the C- to the N-terminal face of MCM, which will form the translocation strand upon replisome activation (Figure 2c). The h2i pore loops more loosely follow a staircase pattern and engage both strands of the double helix (ED Fig. 4).

The zinc fingers of the N-terminal B domain form a 13 Å passage that is too narrow to accommodate B-form DNA where the two MCM rings dimerise (Fig. 2a). For the two DNA strands to traverse this tight pore, 5 bp of the double helix become underwound and one bp is broken (Fig. 2a). The resulting orphan bases are each stabilised symmetrically by MCM5 R195 through a cation-pi interaction, while L209 engages the neighbouring base through a hydrophobic contact that compensates for the missing base stacking (Fig. 2d). In this configuration each MCM hexamer is engaged to the orphan base of what will become the translocation strand (the leading strand) once the replisome is activated. The close match between the structure of hDH from cells^4^ and our hDH assembled in vitro indicates that hDH loading alone is sufficient to untwist DNA and break base pairing, with no additional activation steps required.

To assess the role of R195 and L209 of MCM5 in hDH assembly and bubble formation, we generated MCM complexes containing single and double MCM5 mutants. Fig. 2e shows that the individual single mutants as well as the double mutant loaded MCM almost as well as the wild type proteins. The DH assembled with the R195A, L209G double mutant is structurally very similar to the wild type hDH (ED Fig. 5). Notably, DNA engagement by the MCM motor domain is identical to wild type and the DNA between hexamers is still distorted and untwisted to the same degree, although no breakage of DNA base-pairing could be resolved (Fig. 2f). Thus, R195 and L209 are not required for DNA untwisting, but rather, they stabilise the orphan DNA bases formed upon untwisting.

We noticed that the DNA protected after 0.5 min Benzonase digestion was slightly less intense and smaller with the mutants than with the wild type proteins despite the structural similarity of wild type and mutant hDH and DNA within the hDH central channels (Fig. 2g). We considered that the smaller size might indicate that the mutant hDH is more mobile on DNA than the wild type: movement of hDH during Benzonase digestion might expose DNA that was within the central channel at the start of the digestion resulting in a shorter product.

Consistent with this, the DH footprint generated by the human DH was stable with increasing time of digestion up to 20 min whilst the human AG double mutant footprint disappeared with time and was gone by 20 min This is not because the AG DH is less stably bound to DNA than wild type, because the footprint is still detected after 20 min preincubation before adding benzonase for 0.5 min Therefore, the MCM5 AG mutant can efficiently assemble a DH containing unwound DNA, but this DH assembled is more mobile on the DNA.

### Human OCCM Related Complexes

To begin to examine the mechanism of human DH assembly, we used the poorly hydrolysable ATPγS analogue of ATP. In budding yeast, ATPγS supports the recruitment of the ORC-CDC6-CDT1-MCM (OCCM) complex^6,26,27^, an early intermediate in DH assembly^28^. In the Benzonase footprinting assay, yeast OCCM generates a protection of around 50bp consistent with the length of DNA within the ORC-CDC6 and MCM central channels^29^ (ED Fig. 6a); a footprint of a similar size was seen with human proteins in ATPγS (Fig. 3a), with an additional, fainter band of ∼120 bp. Formation of these products with truncated human proteins required ORC1-5, CDT1 and MCM, but did not require ORC6; surprisingly, in the absence of CDC6, both bands were also visible, though at much reduced levels.

**Figure 3.**
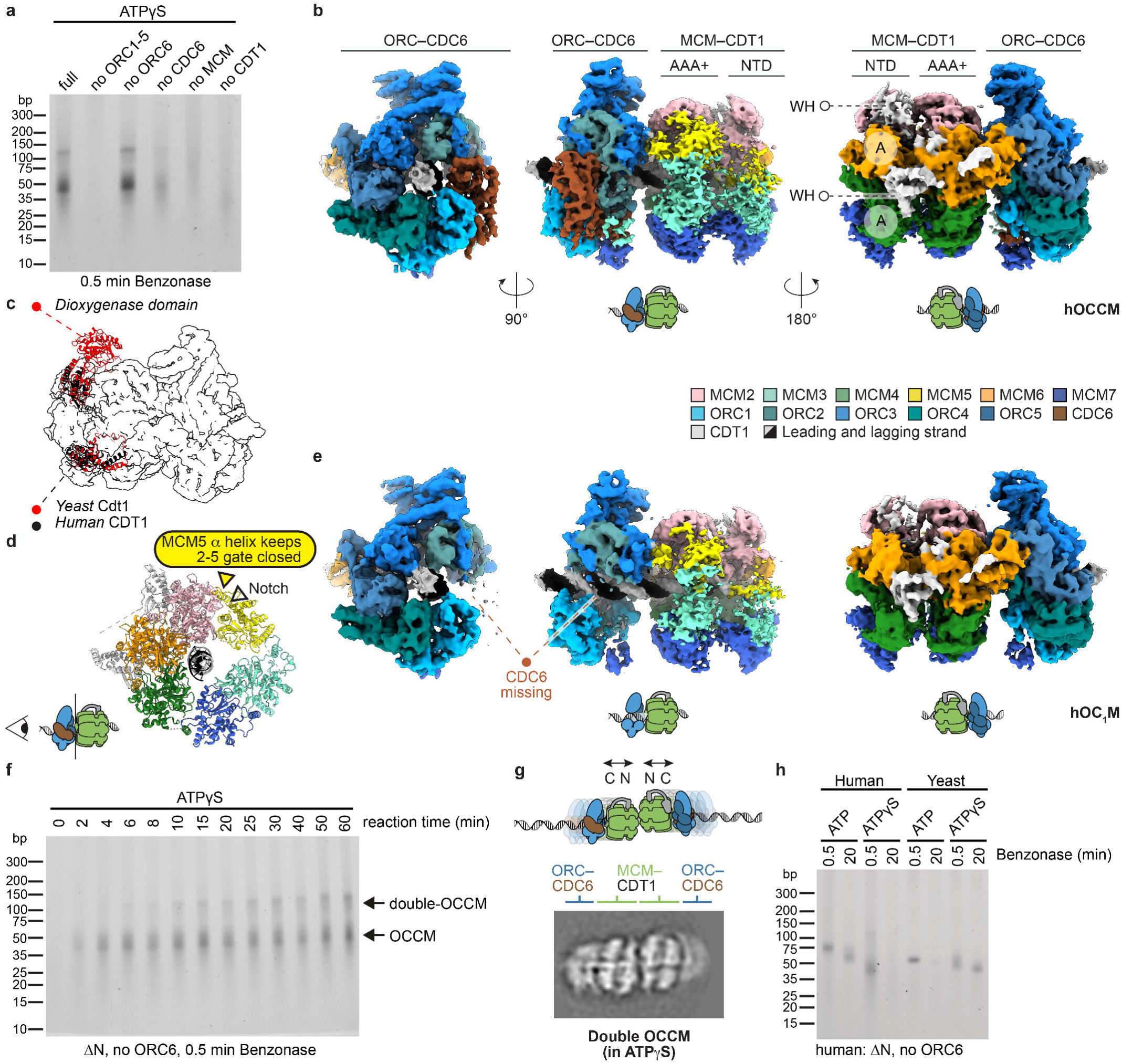
Human OCCM complexes. **a**, Nuclease footprinting assay performed with ATPγS. Proteins were omitted as indicated. **b**, Three views of the hOCCM structure. The WH domains of CDT1 (white) interdigitate between the A domain of MCM6 and MCM4. **c**, Yeast Cdt1 (red) and human CDT1 (black) establish the same interactions with the MCM complex in OCCM. **d**, The MCM2-5 gate is notched but topologically closed by an MCM5 alpha helix packing against the MCM2 ATPase. **e**, Three views of the hOC1M structure. **f**, Reaction timecourse in ATPγS. **g**, Cryo-EM 2D average of a double OCCM. **h**, Nuclease footprinting assay with human and yeast MCM loading reactions using either ATP or ATPγS. The experiment was performed at 30°C.

To characterise the structure stabilised by ATPγS in reconstituted human MCM recruitment onto nucleosome-capped ARS1 DNA, we analysed the reaction by cryo-EM. We solved a 3.8 Å resolution structure of a complex where ORC1-5, CDC6, MCM and CDT1 could be recognised (human OCCM, or hOCCM, Fig. 3b and ED Fig. 6b-f and ED Fig. 7). As in yeast OCCM, the motor domain of MCM is oriented towards the C-terminal face of a hexameric ORC1-5/CDC6 ring. The C-terminal winged-helix (WH) domains of MCM3, 6 and 7 contact the ORC5-4, ORC1-CDC6 and CDC6-ORC2 interfaces respectively. Notably, the MCM4 WH domain that engages ORC in yOCCM^29^ is invisible in the human structure (ED Fig. 8a-c). Conversely, similar to yOCCM, the human ORC-CDC6 and MCM rings are aligned so that duplex DNA can traverse the entire length of the complex. While the cryo-EM density is of sufficient quality to only build 39 bp, we note that OCCM central channel is 185 Å long, which is enough to protect the ∼50 bp measured with Benzonase footprinting. Inside the channel, ORC2, ORC4 and CDC6 touch both DNA strands, and MCM4,6,2 track along the leading strand, via PS1 pore loops arranged in a staircase configuration (ED Fig. 4).

ORC6 is absent in the human OCCM structure, which is not surprising given that omitting ORC6 has no effect on OCCM formation in ATPγS according to DNA footprinting (Fig. 3a), and 2D averages obtained in these conditions appear indistinguishable from reactions where ORC6 is included (ED Fig. 2b,d). Human CDT1 is shorter than the yeast counterpart, as it lacks the N-terminal dioxygenase domain that forms a handle-like feature in the yeast complex^29^ (Fig. 3c and ED Figure 8a). Two tandem winged helix domains of CDT1 interdigitate between the MCM6-2 and MCM4-6 “A” domains, respectively. The two CDT1 WH domains are bridged by an alpha helical element that packs against the MCM6 WH domain, connecting the N and C-terminal tiers of the MCM ring along its outer surface (Fig. 3b). Taken together, CDT1 incorporation into in the hOCCM complex matches observations made for the yOCCM^29^. This is unexpected given that previous co-expression and reconstitution attempts failed to detect any interaction between CDT1 and MCM, which instead form a stable intermediate in the yeast MCM loading reaction. As for the yeast complex^29^, the MCM3 and MCM5 subunits are poorly resolved in the OCCM structure and in the N-terminal side of MCM in particular (Fig. 3b). The MCM2-5 ATPase gate is cracked yet topologically closed, with a recognisable alpha helical element from MCM5 packed against the MCM2 ATPase domain, keeping the subunit interface sealed (Fig. 3d).

A subcomplex of OCCM was solved to 4.1 Å resolution, which contains ORC, CDT1 and MCM but lacks CDC6 (OC1M, Fig. 3e). Compared to the full hOCCM complex, ORC and MCM are bridged by MCM6 and MCM7 WH domains but not MCM3 WH domain, which cannot be averaged when CDC6 is absent (ED Fig. 8b,c). The ORC2 WH domain also becomes invisible when CDC6 is absent, exposing a segment of B-form DNA clamped by ORC1-5 within the OCCM core (Fig. 3e). The OC1M complex could either be a product of OCCM disassembly or form idependently of CDC6 recruitment. Despite the weaker signal, the OCCM-like Benzonase footprint observed when CDC6 is omitted support the latter scenario, as do the OCCM-like 2D class averages obtained by negative stain EM (Fig. 3a, ED Fig. 2f).

The OCCM-related structures described so far explain the ∼50 bp but not the ∼120 bp benzonase footprint band. We noted that the ∼50 bp OCCM band appeared within 2-4 min; however, the larger ∼120 bp band accumulated much more slowly and didn’t reach a maximum until 30-60 min of incubation (Fig. 3f). Geminin, when added after 9 min, led to the disruption of the ∼50 bp band but not the ∼120 bp band (ED Fig. 8d). The larger footprint was roughly double the size of the OCCM, and we considered that it might be two adjoined OCCM complexes. Consistent with this, head-to-head double OCCM class averages were seen in ATPγS reactions after 45 min (Fig. 3g). In these double OCCM 2D averages, the MCM N-termini are closely associated and well resolved, indicating a uniform subunit register. One possible explanation for how the double OCCM forms is that it arises from the encounter of mobile single OCCM assemblies. Consistent with the idea that the OCCM is mobile, Fig. 3h shows that, whilst the human DH footprint was still seen after 20 min digestion, the human OCCM footprint disappeared by 20 min digestion. By contrast, the yDH footprint had disappeared after 20 min of Benzonase digestion, whilst the yeast OCCM footprint was still there after 20 min digestion. Taken together, these results indicate that hDH is immobile and hOCCM is mobile; by contrast, yDH is mobile and yOCCM is immobile.

### Fully loaded single MCM hexamers

Two fully loaded MCM single-hexamer species were identified in both the recruitment (ATPγS) and loading (ATP) datasets. One species is the isolated single hexamer (hSH) topologically bound to duplex DNA, which we solved to 3.2 Å resolution from the ATP dataset, yielding a map of sufficient quality to distinguish ADP from ATP within the catalytic centres (Fig. 4a, ED Fig. 9a-c and ED Fig. 10a). The 3.4 Å structure obtained from ATPγS data (ED Fig. 7) presents no obvious difference and is not further discussed. Unlike for the OCCM complex, both the N- and the C-terminal tiers are well-resolved and the MCM2-5 gate is completely shut. The DNA grip of the PS1 pore loops matches that observed for the hDH, with MCM7,4,6 engaging the leading strand template (Fig. 4b). The path of the double helix is less distorted compared to the hDH structure, which positions L209 too far to contact the double helix and R195 retracted and engaged to a DNA-backbone phosphate. As a result, no DNA untwisting nor melting can be observed in the hSH structure (Fig. 4c). Despite this difference, nucleotide occupancy matches that of the hDH, with one apo (MCM7-4), one ATP and all other sites bound to ADP (Fig. 4a).

**Figure 4.**
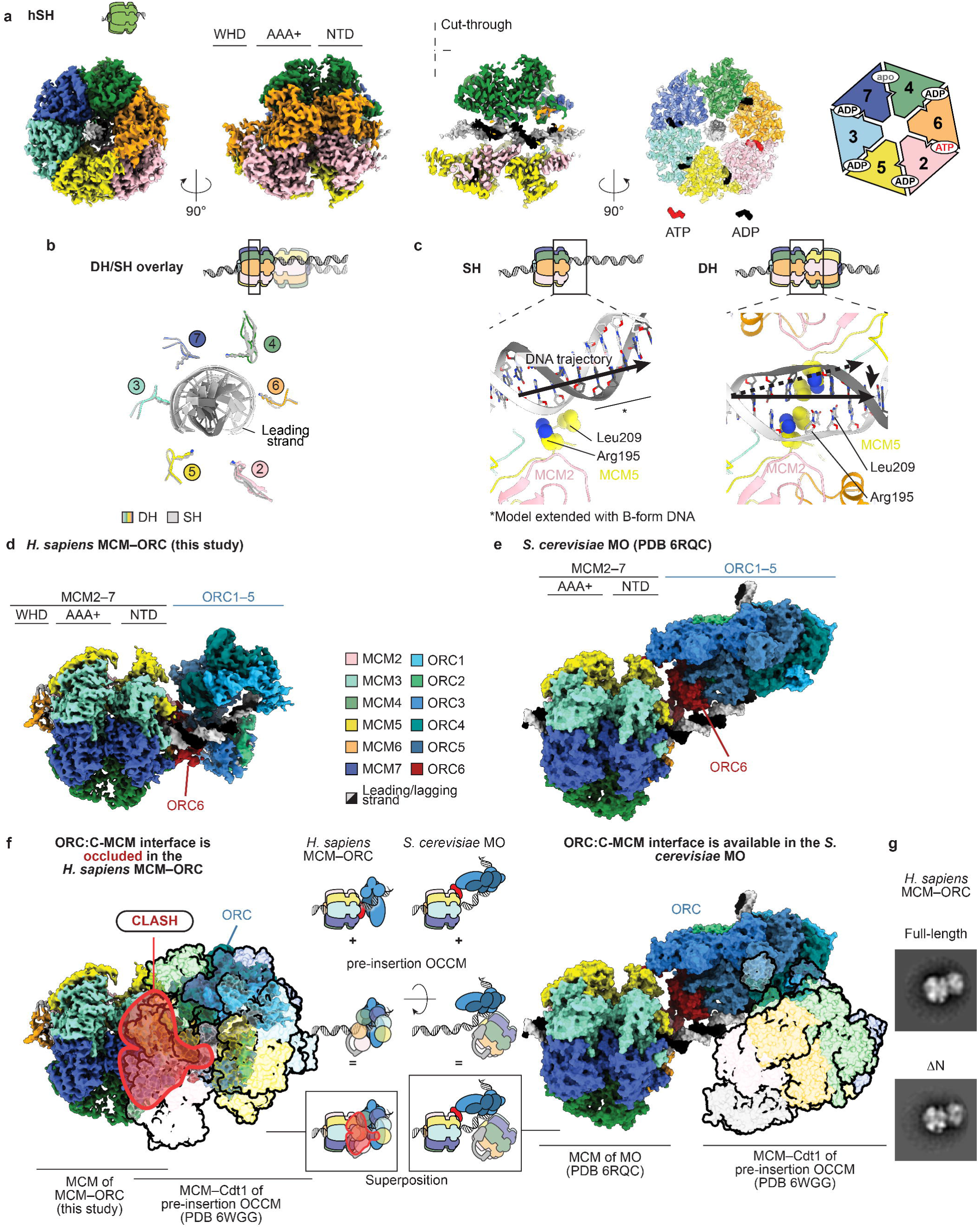
Single-loaded MCM complexes. **a**, Surface rendering and cut-through view of a hSH loaded onto duplex DNA. The C-terminal ATPase site is shown with nucleotides bound at subunit interfaces. **b**, A superposition of PS1 pore loops and DNA extracted from hDH and hSH show the same configuration. **c**, MCM hexamer dimerisation changes the path of duplex DNA leading to untwisting and opening of the double helix. **d**, Surface rendering of the hMO* structure. **e**, Surface rendering of the yMO structure highlights a different configuration of the ORC complex compared to hMO*. ORC6 is bridges N-terminal MCM and ORC1-5 in both structures. **f**, A steric clash obtained by modelling the pre-insertion yOCCM (PDB entry 6WGG) demonstrates that hMO* cannot support recruitment of a second MCM, unlike what was observed for the yMO complex. **g**, Truncated and full length ORC1, CDC6 and CDT1 yield the same hMO* complex, as observed by negative stain 2D averaging.

A second assembly contains a fully loaded hSH bound to ORC (Fig. 4d, ED Fig. 9d-f and ED Fig. 7). In this structure, ORC1-5 (solved to 4 Å resolution by local refinement, ED Fig. 9g,h) binds and bends DNA. Within the MCM ring (solved to 3.5 Å resolution as a separate body ED Fig. 9i,j), the DNA grip appears indistinguishable from that of the isolated hSH. ORC6 is essential for the formation of this intermediate (ED Fig. 2) and bridges between ORC1-5 and the N-terminal side of MCM. Here, the N-terminal cyclin box domain of ORC6 is sandwiched between ORC4 and the N-terminal Zinc Finger (ZnF) domains of MCM2 and MCM6. The C-terminal cyclin box domain of ORC6 instead connects ORC5 with the ZnF domains of MCM4 and MCM6 (ED Fig. 10b). A human specific IDR in ORC6 would make it physically possible for a conserved C-terminal alpha helix of ORC6 to contact ORC3 as previously observed in yeast^28,29^ and fly^30^. Due to limited local resolution, however, we cannot comment on whether this interaction exists in the human complex (ED Fig. 10b). Given that the C-terminal alpha helix is essential for ORC6 recruitment to ORC in lower^28,29^ and higher eukaryotes^30^, we consider it likely that this interaction exists in the ORC6-dependent human MCM-ORC.

Time-resolved cryo-EM and single-molecule work with yeast proteins showed that ORC sequentially binds first the C- (in OCCM) and then the N-terminal side of MCM (in the MCM-ORC, yMO, loading intermediate, Fig. 4e). In yMO, Orc6 straddles across the closed Mcm2-5 gate (ED Fig. 10a) of a ySH and orients ORC so that it can load a second ySH to form a yDH (Figure 4f). In the human structure, the N-terminal cyclin box of ORC6 contacts MCM2 as seen in the yMO complex. The N-terminal ORC6 engagement with ORC4 is only seen in the human complex and not in yMO. Also consolidated by other human-specific interactions (ED Fig. 10b), this ORC6-ORC4 contact collapses the structure in a folded configuration where recruitment of a second hSH is sterically impeded (Fig. 4f). To mark the structural differences with yMO, we named the human MCM-ORC complex, hMO*.

## Discussion

Although yeast and humans use ORC, CDC6 and CDT1 to load an MCM DH around double stranded DNA, our work shows that there are clear differences in how the two organisms accomplish this feat. For example, CDT1 is pre-bound to MCM in yeast where it stabilises an open form of the MCM hexamer whilst human CDT1 does not associate with free MCM but instead joins the complex during MCM recruitment. Despite this difference, the interactions of CDT1 with MCM are very similar in hOCCM and yOCCM. In yeast, the initial recruitment of MCM to origins is dependent upon interaction between CDC6 and a WH domain at the C-terminus of MCM3^5^; this interaction is conserved in hOCCM but CDC6 is not essential for MCM recruitment and interactions of ORC with WH domains in MCM6 and MCM7 stabilise the OC1M structure in the absence of CDC6.

We see no MCM loading without ORC1-5 or CDC6, consistent with genetic analysis indicating that they are essential for human cell viability^31^ though we cannot yet rule out the possibility that individual subunits of ORC1-5 may be dispensible for hDH loading^32,33^.

ORC6 is essential for yDH loading, but our results show that hDH is loaded *in vitro* via at least two distinct pathways, one ORC6-dependent and one ORC6-independent (Fig. 5). It is unclear whether both pathways are used *in vivo*: on one hand a Meier-Gorlin Syndrome mutant mapping to the C-terminal alpha-helix of ORC6 that interacts with ORC3 and inhibits ORC6 interaction with ORC1-5 also inhibits MCM loading in *Drosophila melanogaster*^34^; on the other hand, depletion of ORC6 from human U2OS cells was shown to have no effect on MCM loading levels in a single cell cycle^35^.

**Figure 5.**
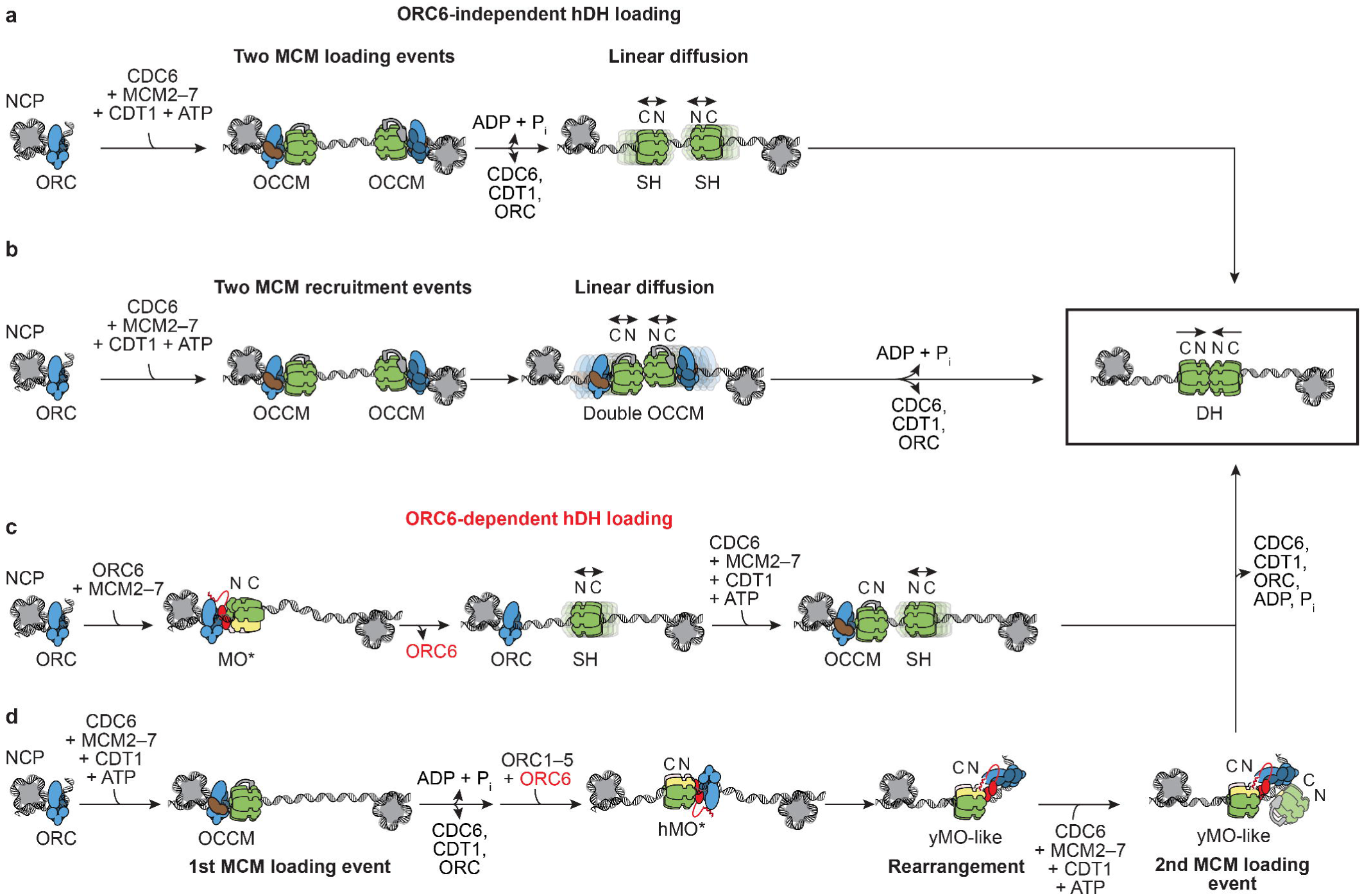
Mechanisms of hDH loading. **a**, MCM can be loaded via an ORC6 independent pathway, via two inverted hOCCM complexes that load two hSHs in a process that requires ATP hydrolysis. Free diffusion along duplex DNA would then lead to hDH formation. **b,** In a variation of the same mechanism, two hOCCMs assembled around duplex DNA are free to diffuse and form a dOCCM. ATP hydrolysis then promotes hDH formation. **c,** MCM loading with full-length ORC1 can occur in an ORC6 dependent manner and might go through the hMO* intermediate. hMO* might recruit a hSH through an ORC6 interaction with N-terminal MCM. Following hSH release, the same ORC could recruit a second hSH via C-terminal MCM interaction (hOCCM intermediate), resulting in two N-terminally facing hSHs that can assemble a hDH. **d,** Alternatively, a first hSH could be loaded via OCCM. ORC6 would then interact with the N-terminal domain of hSH forming hMO*. A structural change would then occur, which causes hMO* to transition to a yMO-like state, where hORC is competent for the recruitment of a second hSH, eventually leading to hDH formation. We have however not observed a yMO-like complex formed with human proteins.

ORC6-dependent loading may occur through formation of the ORC6-dependent hMO* complex. hMO* contains a closed-ring, loaded SH with its N-terminal domain bound to ORC. hMO* was also seen in the absence of CDT1 and CDC6, suggesting that the SH in hMO* may not arise from OCCM but may instead be loaded directly by ORC alone. ORC6 in hMO* engages the N-terminal domains of MCM2,4 and 6; this is the same side of MCM bound by CDT1 in the yMCM-CDT1 loading intermediate (ED Fig. 10), which promotes loading of a single MCM hexamer in yeast. Loading of hDH, however, still requires CDC6 and CDT1, arguing that at least one SH in the DH must be loaded via OCCM. The yeast SH has a short lifetime on DNA^9^, so it is possible that the role of ORC in hMO* is to stabilise SH on DNA. The position of ORC at the N-terminus of the SH in hMO*, however, would block the recruitment of a second hexamer as seen in yMO (Fig. 4f) and block hDH assembly via a second SH. Therefore, for hMO* to be a fruitful intermediate, ORC must somehow release its grip on MCM. Because of how the N- and C-terminal cyclin boxes are connected in ORC6, human MCM and ORC cannot recapitulate the exact same architecture seen in yMO. The hMO* structure could however transition from the collapsed state described here to a more open configuration similar to yMO, which we have not been able to observe. The ORC1 IDR is critical for ORC6-dependent loading but is not required for hMO* formation, and could have a role in this hMO* rearrangement, creating the space for ORC-mediated recruitment of a second hSH. Alternatively ORC1 IDR could act on hMO* by disengaging ORC from MCM, perhaps prompted by interaction with a second, correctly oriented SH loaded from OCCM. Such a role for the ORC1 IDR could also explain why ORC6 inhibits MCM loading with truncated proteins: in these reactions hMO* is a dead-end complex that cannot be converted to hDH because ORC cannot release from hMO*.

ORC6-independent hDH loading may occur via two oppositely oriented and independently loaded single hexamers. This is fundamentally similar to assembly of yDH from two high affinity binding sites when yMO formation is blocked, for example, by CDK phosphorylation of ORC2 [ref ^9^] except that there are no specific sequences to correctly orient hexamer loading. This might appear wasteful since only 25% of randomly loaded MCM pairs would be predicted to be in the correct orientations to assemble DH. Nonetheless, hDH assembly *in vitro* by ORC containing truncated ORC1 and lacking ORC6 is quite efficient; amounts of hDH assembled are similar to amounts of yDH assembled from specific DNA sequences under similar conditions (e.g. Fig. 1x). A second possible mechanism captitalises on the fact that human ORC is not a sequence specific DNA binding protein: because human ORC is not anchored at a specific sequence, intermediates such as OCCM are not static, in contrast to yeast, but rather can diffuse on DNA. Loading of hDH via a double OCCM could, for example, guarantee correct orientation of two hexamers before loading. The ORC6-independent pathway is inefficient with wild type, full length ORC1. This may indicate that the role of the ORC1 IDR is to prevent ORC6-independent hDH loading. Alternatively, some post-translational modification may counteract ORC1 IDR inhibition, allowing ORC6-independent loading to occur in a regulated manner.

Although the structures of hDH and yDH are very similar, their engagement with DNA has two major differences. Firstly, hDH has a 5bp bubble of unwound DNA between the hexamers, stabilised by R195 and L209 of MCM5. R195 and L209 are not required for unwinding, but stabilise the bubble. hDH containing two hexamers with the R195, L209 mutations could not be isolated from chromatin when expressed in cells suggesting that bubble formation may be required for hDH assembly^4^. Our results indicate that this mutant can form hDH, but the hDH formed is mobile on DNA. Thus, the inability to detect the mutant in cells may reflect the loss of bound DNA after Benzonase treatment during purification, which might render the mutant hDH unstable. The second major difference is that DNA engagement by yDH within the central channel is displaced by approximately 5 bp relative to hDH. In comparing the DNA grip of various MCM structures, yDH emerges as an outlier: ySH, hSH and hMO* all grip the DNA very similar to hDH. DNA slippage within the central channel during yDH loading might dissipate torsional strain. DNA in the yMO is already in the same configuration as yDH indicating that this slippage can happen before the final steps in yDH formation. In yeast, origin melting begins within the AAA+ domains of each hexamer during helicase activation, not at the interface between hexamers^36^. It will be interesting to see where melting begins during human helicase activation.

Our results show that hDH can be assembled efficiently in the absence of both human origin sequences and nucleosomes. Thus, any nucleosome-free stretch of DNA of appropriate length in the genome may be sufficient for hDH assembly and origin function *in vivo*.

Efficient assembly of an hDH that is virtually identical to the hDH isolated from cells represents a first step toward reconstitution of DNA replication with purified human proteins.

## Materials and Methods

### Protein sequences

Human cDNAs were cloned from RPE-1 and U2-OS cells by total RNA isolation, oligo-dT reverse transcription, followed by PCR with gene-specific oligonucleotides and cloning into expression vectors. Sanger sequencing showed that the cloned cDNA coded for the reference protein sequence for the following proteins: ORC1, ORC2, ORC3(NCBI: iso2, Uniprot: canonical iso), ORC4, ORC5, ORC6, MCM2, MCM3, MCM5, MCM6, MCM7, CDC6, GMNN. For MCM4 and CDT1 the cloned cDNA coded for a natural variant that differed from NCBI reference protein sequences but according to the Genome Aggregation Database (gnomad.broadinstitute.org) represented the majority allele (allele frequency > 0.5): MCM4_L650M (allele frequency: 0.862), CDT1_C234R (allele frequency: 0.999). These alleles were used and considered wild type throughout this study.

### Expression and purification of human MCM2-7

MCM2-7 was expressed in insect cells using the biGBac baculovirus expression vector system^10^. Human cDNAs of MCM2, MCM3, MCM4, MCM5, MCM6, and MCM7 were cloned into pLIB. MCM3 was subcloned to contain an N-terminal TEV protease-cleavable Flag tag. pLIB-derived expression cassettes of MCM4, MCM5, MCM6, MCM7 were subcloned into pBIG1a (pBIG1a:MCM4,5,6,7) and expression cassettes of MCM2 and Flag-MCM3 were subcloned into pBIG1b (pBIG1b:MCM2,Flag-3). Expression cassettes of these two vectors were subcloned into pBIG2ab (pBIG2ab:MCM4,5,6,7,2,Flag-3). Baculovirus was generated and amplified in Sf9 cells (Thermo Fisher, 12659017) using the EMBacY baculoviral genome^37^. For protein expression, Sf9 cells were infected and collected 52 hours after infection, flash-frozen and stored at -80°C.

Cell pellets were thawed on ice in MCM buffer (50 mM HEPES/KOH pH 7.6, 100 mM potassium glutamate, 5 mM magnesium acetate, 10% glycerol, 0.02% NP-40, 1 mM DTT) + protease inhibitors (1 tablet / 50 ml Roche Complete Ultra EDTA-free, 10 µg/ml leupeptin, 10 µg/ml pepstatin A, 1 mM AEBSF, 1 µg/ml aprotinin, 2 mM benzamidine). Cells were resuspended and lysed by Dounce homogenisation using a tight-fitting pestle. The lysate was cleared by centrifugation (158,000 g, 4°C, 1 h) and incubated with anti-FLAG M2 affinity gel (Sigma-Aldrich, A2220) for 1 h at 4°C with rotation. The beads were transferred to a Econo-Pac gravity flow column (Bio-Rad, 732-1010) and washed with 2x 20 column volumes (cv) MCM buffer, 1x 20 cv MCM buffer + 5 mM ATP (Sigma-Aldrich, A2383). For dephosphorylation, the beads were resuspended in MCM buffer + 0.2 mg/ml lambda protein phosphatase, 1 mM manganese(II) chloride and incubated for 1 h at 4°C. The beads were washed with 2x 20 cv MCM buffer. For elution and proteolytic tag removal the beads were resuspended in 5 cv MCM buffer + 80 µg/ml TEV protease, 0.1 mg/ml 3xFlag peptide for 2 h at 4°C. The eluate was concentrated using an Amicon Ultra-15 concentrator (Merck, UFC903024) and further purified by gel filtration using a HiLoad 16/600 Superdex 200 pg column equilibrated in MCM buffer. Fractions containing stoichiometric MCM2-7 were concentrated, flash frozen in liquid nitrogen, and stored at -80°C.

### Expression and purification of human ORC1-5

ORC1-5 was expressed in baculovirus-infected insect cells. The coding sequences of human ORC1, ORC2, ORC3, ORC4 and ORC5 were codon-optimised for *S. frugiperda*, synthesised (GeneArt, Thermo Fisher Scientific) and subcloned into modified pBIG1 vectors that contain a pLIB derived polyhedrin expression cassette. ORC1 was subcloned into pBIG1a with a TEV protease-cleavable N-terminal 3xFlag tag. ORC2 was subcloned into pBIG1b, ORC3 into pBIG1c, ORC4 into pBIG1d, ORC5 into pBIG1e. Expression cassettes from these five vectors were subcloned into pBIG2abcde (pBIG2abcde:Flag-ORC1,2,3,4,5). Baculoviruses were generated using EMBacY and Sf9 cells. To express ORC1-5, Sf9 cells were infected, collected 52 hours after infection, flash-frozen and stored at -80°C.

Cell pellets were thawed in ORC1-5 lysis buffer (50 mM HEPES/KOH 7.6, 650 mM potassium chloride, 5 mM magnesium acetate, 1 mM ATP, 10% glycerol, 0.02% NP-40, 1 mM DTT, 2 mM benzamidine) + protease inhibitors (1 tablet / 50 ml Roche Complete Ultra EDTA-free, 10 µg/ml leupeptin, 10 µg/ml pepstatin A, 1 mM AEBSF, 1 µg/ml aprotinin). Cells were lysed by Dounce homogenisation, and the lysate cleared by centrifugation (158,000 g, 4°C, 1 h). ORC1-5 was bound to anti-FLAG M2 affinity gel for 1 h at 4°C. The column was washed with 2x 20 cv ORC1-5 lysis buffer + 4 mM ATP. For dephosphorylation, the beads were resuspended in 1 cv ORC1-5 lysis buffer + 0.2 mg/ml lambda protein phosphatase, 1 mM manganese(II) chloride, potassium chloride adjusted to 650 mM, and incubated at 4°C for 1 h. The column was washed with ORC1-5 lysis buffer. For elution and proteolytic tag removal, the beads were resuspended in 5 cv ORC1-5 lysis buffer + 80 µg/ml TEV protease,

0.1 mg/ml 3xFlag peptide, potassium chloride adjusted to 650 mM and were incubated at 4°C for 2 h. To remove TEV protease, the eluate was supplemented with 35 mM imidazole pH 8.0 and incubated with Ni-NTA Agarose (Invitrogen, R90115) for 1 h at 4°C. The flowthrough was concentrated and further purified by gel filtration using a HiLoad 16/600 Superdex 200 pg column (Cytiva) equilibrated in ORC1-5 SEC buffer (50 mM HEPES/KOH pH 7.6, 650 mM potassium chloride, 5 mM magnesium acetate, 10% glycerol, 0.02% NP-40, 1 mM DTT). Fractions containing stoichiometric ORC1-5 were concentrated, flash-frozen, and stored at -80°C.

### Expression and purification of human CDC6

CDC6 was expressed in insect cells. Human cDNA of CDC6 was cloned into pLIB to contain an N-terminal TEV protease-cleavable Flag tag. The baculovirus was generated in Sf9 cells using the EMBacY genome. For expression, Sf9 cells were infected and the culture harvested 52 hours after infection. The cell pellet was flash-frozen in liquid nitrogen and stored at -80°C.

The cell pellet was thawed on ice in CDC6 lysis buffer (50 mM HEPES/KOH pH 7.6, 650 mM potassium chloride, 5 mM magnesium acetate, 4 mM benzamidine, 1 mM ATP, 10% glycerol, 0.02% NP-40, 1 mM DTT) + protease inhibitors (1 tablet / 50 ml Roche Complete Ultra EDTA-free, 10 µg/ml leupeptin, 10 µg/ml pepstatin A, 1 mM AEBSF, 1 µg/ml aprotinin). Cells were lysed with a Dounce homogeniser and the lysates centrifuged (158,000 g, 4°C, 1 h). The cleared lysate was incubated with anti-FLAG M2 affinity gel for 1 h at 4°C. The beads were washed with 2x 20 cv CDC6 lysis buffer + 4 mM ATP. For dephosphorylation, the beads were resuspended in CDC6 lysis buffer + 0.2 mg/ml lambda protein phosphatase, 1 mM manganese(II) chloride and incubated for 1 h at 4°C. The column was washed with CDC6 lysis buffer, followed byCDC6 HTP-wash buffer (50 mM potassium phosphate pH 7.6, 75 mM potassium acetate, 5 mM magnesium acetate, 0.1% Triton X-100, 1 mM DTT, 2 mM ATP). For elution, the beads were resuspended in CDC6 HTP-wash buffer + 80 µg/ml TEV protease, 0.1 mg/ml 3xFlag peptide and incubated for 2 h at 4°C. To remove TEV protease, the eluate was supplemented with 30 mM imidazole pH 8.0 and incubated with Ni-NTA agarose for 1 h at 4 °C. The CDC6-containing flowthrough was collected. A hydroxyapatite column was prepared by resuspending 2 g Bio-Gel HTP Hydroxyapatite (Bio-Rad, 130-0420) in 12 ml CDC6 HTP-wash buffer. To remove fine particles, the beads were allowed to settle for 2 min and the supernatant was removed. Two more times, the beads were resuspended in 10 ml CDC6 HTP-wash buffer, allowed to settle for 2 min, and the supernatant removed.

Then 4 ml of a 50% slurry were incubated with CDC6 for 15 min at 4°C. The beads were transferred to a gravity flow column and washed with 2 ml CDC6 HTP-wash buffer, followed by 5 ml CDC6 HTP-rinse buffer (50 mM potassium phosphate pH 7.6, 150 mM potassium acetate, 5 mM magnesium acetate, 0.1% Triton X-100, 15% glycerol, 1 mM DTT). CDC6 was eluted by applying 10 ml CDC6 HTP-elution buffer (50 mM potassium phosphate pH 7.6, 400 mM potassium acetate, 5 mM magnesium acetate, 0.1% Triton X-100, 15% glycerol, 1 mM DTT). The eluate was dialysed 2x 1 h at 4°C against CDC6 dialysis buffer (50 mM HEPES/KOH pH 7.6, 650 mM potassium chloride, 5 mM magnesium acetate, 10% glycerol, 0.02% NP-40, 1 mM DTT). CDC6 was concentrated, flash-frozen, and stored at -80°C.

### Expression and purification of human CDT1

CDT1 was expressed in insect cells. The coding sequence of human CDT1 was codon-optimised for *S. frugiperda*, synthesised (GeneArt, Thermo Fisher Scientific) and subcloned into pLIB as a fusion protein with an N-terminal Flag-His-SumoEu1 fusion^38^. The baculovirus was generated in Sf9 cells using the EMBacY genome. Expression cultures were harvested 52 h after infection, snap-frozen in liquid nitrogen and stored at -80°C.

The cell pellet was thawed in CDT1 buffer (50 mM HEPES pH 7.6, 650 mM potassium chloride, 5 mM magnesium acetate, 10% glycerol, 0.02% NP-40, 1 mM DTT) + protease inhibitors (1 tablet / 50 ml Roche Complete Ultra EDTA-free, 10 µg/ml leupeptin, 10 µg/ml pepstatin A, 1 mM AEBSF, 1 µg/ml aprotinin). Cells were lysed using a Dounce homogeniser. The lysate was cleared by centrifugation (158,000 g, 4°C, 1 h) and incubated with anti-Flag M2 affinity gel for 1 h at 4°C. The beads were transferred to a gravity flow column and washed twice with CDT1 buffer + 5 mM ATP. The beads were resuspended in CDT1 buffer + 0.2 mg/ml lambda protein phosphatase, 1 mM manganese(II) chloride and incubated for 1 h at 4°C. The beads were washed with CDT1 buffer. For proteolytic elution the beads were resuspended in CDT1 buffer + 80 µg/ml His-SENP1_EuH protease^38^ and incubated for 2 h at 4°C. To remove His-SENP1_EuH protease, 35 mM imidazole pH 8.0 was added and the eluate incubated with Ni-NTA agarose for 1 h at 4°C. The CDT1-containing flowthrough was concentrated and further purified by gel filtration on a Superdex 200 Increase 10/300GL column (Cytiva) using CDT1 buffer. CDT1 containing fractions were concentrated, snap-frozen, and stored at -80°C.

### Expression and purification of human ORC6

ORC6 was expressed in *E. coli*. The coding sequence of human ORC6 was codon-optimised for E. coli, synthesised (GeneArt, Thermo Fisher Scientific) and subcloned into pK27Sumo to encode the fusion protein His-Sumo-ORC6. The protein was expressed using the strain T7express lysY (NEB, C3010I). Expression was induced with 0.4 mM IPTG at OD600 ∼0.6 at 16°C. The culture was harvested after 16 h, flash-frozen, and stored at -80°C.

The cell pellet was thawed in ORC6 lysis buffer (50 mM HEPES pH 7.6, 500 mM potassium chloride, 5 mM magnesium acetate, 10% glycerol, 0.02% NP-40, 1 mM DTT, 35 mM imidazole) + protease inhibitors (1 tablet / 50 ml Roche Complete EDTA-free, 1 mM AEBSF), lysozyme. The cells were resuspended and lysed by sonication. The lysate was centrifuged (158,000 g, 4 °C, 1 h) and the cleared lysate incubated with Ni-NTA agarose (Invitrogen) for 1 h at 4°C. The beads were transferred to a gravity flow column and washed 2x with ORC6 lysis buffer + 5 mM ATP and 2x with ORC6 lysis buffer. The protein was eluted with ORC6 elution buffer (50 mM HEPES/KOH pH7.6, 200 mM potassium chloride, 5 mM magnesium acetate, 10% glycerol, 0.02% NP-40, 1 mM DTT, 300 mM imidazole pH 8.0). The Sumo-specific protease His-Ulp1 was added at 0.08 mg/ml and the protein was dialysed overnight against ORC6 SEC buffer (50 mM HEPES/KOH pH7.6, 200 mM potassium chloride, 5 mM magnesium acetate, 10% glycerol, 0.02% NP-40, 1 mM DTT). To remove His-Ulp1 protease and His-Sumo, the imidazole concentration was adjusted to 35 mM and the protein incubated with Ni-NTA agarose (Invitrogen) for 1 h at 4°C. The ORC6 containing flowthrough was concentrated and further purified by gel filtration using a HiLoad 16/600 Superdex 200 pg column (Cytiva) equilibrated in ORC6 SEC buffer. ORC6 containing fractions were pooled, snap-frozen, and stored at -80°C.

### Expression and purification of ORC1-5 1ΔN, CDC6ΔN and CDT1ΔN

The proteins ORC1-5 1ΔN, CDC6ΔN and CDT1ΔN with N-terminal IDR truncations were expressed in insect cells using a Flag-His-SumoEu1 fusion system^38^ and sequences that were codon-optimised for *S. frugiperda* and synthesised (GeneArt, Thermo Fisher Scientific). The SumoEu1 fusions were only partially stable in Sf9 cells giving relatively low yields. Flag-His-SumoEu1-ORC1(391-861) was cloned into pBIG1a containing a pLIB derived polyhedrin expression cassette. The expression cassette was subcloned together with ORC2-5 expression cassettes into pBIG2abcde (pBIG2abcde:Flag-His-SumoEu1-ORC1ΔN,ORC2,3,4,5). Flag-His-SumoEu1-CDC6ΔN(143-560) and Flag-His-SumoEu1-CDT1ΔN(167-546) were cloned into pLIB. Baculoviruses were generated in Sf9 cells using EMBacY. Sf9 expression cultures were collected 52 h after infection, flash-frozen and stored at -80°C.

The three proteins were purified with the same purification protocol with only the gel filtration column differing. Cell pellets were thawed in Wash-300 buffer (50 mM HEPES/KOH pH 7.6, 300 mM potassium chloride, 5 mM magnesium acetate, 1 mM ATP, 10% glycerol, 0.02% NP-40, 1 mM DTT) + protease inhibitors (1 tablet / 50 ml Roche Complete Ultra EDTA-free, 10 µg/ml leupeptin, 10 µg/ml pepstatin A, 1 mM AEBSF, 1 µg/ml aprotinin, 4 mM benzamidine) and lysed using a Dounce homogeniser. The lysate was centrifuged (158,000 g, 4°C, 1h) and the cleared lysate incubated with anti-FLAG M2 affinity gel for 2 h at 4°C. The beads were washed 2x with Wash-300 buffer + 4 mM ATP, and then resuspended in Wash-300 buffer + 0.2 mg/ml lambda protein phosphatase, 1 mM manganese(II) chloride and incubated for 1 h at 4°C. The beads were washed with Wash-300 buffer. For proteolytic elution, the beads were resuspended in Wash-300 buffer + 80 µg/ml His-SENP1_EuH protease^38^ and incubated for 2 h at 4°C. To remove His-SENP1_EuH protease, the eluate was supplemented with 35 mM imidazole pH 8.0 and incubated with Ni-NTA agarose (Invitrogen) for 1 h at 4°C. The flowthrough was concentrated and further purified by gel filtration using SEC-ΔN buffer (50 mM HEPES/KOH pH 7.6, 500 mM potassium glutamate, 5 mM magnesium acetate, 10% glycerol, 0.02% NP-40, 1 mM DTT). ORC1-5 1ΔN was purified using a HiLoad 16/600 Superdex 200 pg column (Cytiva). CDC6ΔN and CDT1ΔN were purified using a Superdex 200 Increase 10/300GL column (Cytiva). Fractions containing the respective protein were concentrated, snap-frozen, and stored at -80°C.

### Expression and purification of human geminin

Geminin was expressed in insect cells. Human cDNA of geminin was cloned into pLIB with an N-terminal Flag tag. The baculovirus was generated in Sf9 cells using EMBacY. Geminin was expressed in Sf9 cells and the culture harvested 52 h after infection. The cell pellet was stored at -80°C.

The cell pellet was thawed in GMNN lysis buffer (50 mM HEPES/KOH pH 7.6, 300 mM potassium chloride, 5 mM magnesium acetate, 10% glycerol, 0.02% NP-40, 1 mM DTT) + protease inhibitors (1 tablet / 50 ml Roche Complete Ultra EDTA-free, 10 µg/ml leupeptin, 10 µg/ml pepstatin A, 1 mM AEBSF, 1 µg/ml aprotinin) and lysed using a Dounce homogeniser. The lysate was cleared by centrifugation (158,000 g, 60 min, 4°C) and incubated with anti-FLAG M2 affinity gel for 1 h. The beads were washed with GMNN lysis buffer + 5 mM ATP, followed by GMNN lysis buffer. Protein was eluted in GMNN lysis buffer + 0.1 mg/ml 3xFlag peptide. The protein was concentrated and further purified by gel filtration on a Superdex 200 Increase 10/300GL column equilibrated in GMNN SEC buffer (50 mM HEPES/KOH pH 7.6, 200 mM potassium chloride, 5 mM magnesium acetate, 10% glycerol, 0.02% NP-40, 1 mM DTT). Geminin containing fractions were concentrated and the protein concentration determined for the homodimer. The protein was flash-frozen, and stored at -80°C.

### Nuclease footprinting assay with human proteins

MCM loading reactions were performed in assay buffer (25 mM HEPES/KOH pH 7.6, 100 mM sodium acetate, 10 mM magnesium acetate, 1 mM DTT). MCM was typically loaded onto a ARS1-containing 10.6 kb plasmid that was purified by caesium chloride density gradient centrifugation (pJY22, ref^39^).

A 20 µl reaction with truncated proteins typically contained 4 nM (27.5 ng/µl) plasmid DNA (10.6 kb), 2 mM ATP, 60 nM MCM2-7, 150 nM ORC6, 150 nM CDC6ΔN, 150 nM CDT1ΔN, and 90 nM ORC1-5(1ΔN). Stocks of MCM2-7, CDC6ΔN, CDT1ΔN, and ORC1-5(1ΔN) were diluted to a 10x working concentration in assay buffer. ORC6 was diluted to a 20x working concentration in assay buffer + 200 mM sodium chloride. Reactions were started by adding MCM2-7, ORC6, CDC6ΔN, CDT1ΔN and ORC1-5(1ΔN) to a mix of DNA and ATP in assay buffer.

A 20 µl reaction with full length proteins typically contained 4 nM (27.5 ng/µl) plasmid DNA (10.6 kb), 2 mM ATP, 60 nM MCM2-7, 150 nM ORC6, 120 nM ORC1-5, 150 nM CDC6, and 150 nM CDT1. MCM2-7 was diluted to 10x working concentration in assay buffer. ORC6 was diluted to 20x working concentration in assay buffer + 200 mM sodium chloride. A 20x OCC mix of ORC1-5, CDC6 and CDT1 was prepared in assay buffer + 650 mM sodium chloride. Reactions were started by adding MCM2-7, ORC6, and the OCC mix to a mix of DNA and ATP in assay buffer.

In experiments that contained truncated proteins as well as their full-length counterparts, ORC1-5 and ORC1-5(1ΔN) were both used at 120 nM and the 20x OCC mix (650 mM sodium chloride) method was used. When proteins were omitted, a salt-containing buffer was added instead to maintain the same salt concentration throughout the experiment.

Reactions were started in 2-min intervals and incubated at 37°C in a thermomixer with shaking at 1,250 rpm. After 30 min of MCM loading, 2 µl of Benzonase nuclease (Sigma-Aldrich, E1014) was added and the mix incubated at 37°C with shaking for either 0.5 min or 20 min depending on the experiment. Then 20 µl of the mix was transferred to a tube containing 10 µl 3x Stop buffer (assay buffer + 100 mM EDTA, 500 µM proteinase K (Sigma-Aldrich, 107393), 1% SDS) and the mix incubated at 37 °C for 20 min with shaking.

The sample was diluted with TE (10 mM Tris/HCl pH 8.0, 1 mM EDTA) to 200 µl and an equal volume of phenol/chloroform/isoamyl alcohol (25/24/1; Invitrogen UltraPure, 15593031) was added. The sample was vortexed for 1 min, transferred to a 5PRIME Phase Lock Gel Heavy spin column (VWR, 733-2478), and spun for 5 min at 20,000 g. The aqueous phase was transferred to a new tube, 20 µl of 3 M sodium acetate pH 5.2, 1 µl of 20 mg/ml glycogen (Thermo Scientific, R0561), and 550 µl ethanol were added and DNA precipitated overnight at -20°C. The DNA was pelleted (20,000 g, 4°C, 40 min), washed with 80% ethanol (20,000 g, 4°C, 10 min), and air-dried. The pellet was resuspended in 3 µl TE. Then 1.5 µl 20% Ficoll 400 (Sigma-Aldrich, F2637) was added, and samples were loaded on a pre-run 4-20% Novex TBE gel (Invitrogen, EC62252BOX). The gel was run using 1xTBE running buffer (Invitrogen, LC6675) at 150 V for 50 min, stained with SYBR Gold (Invitrogen, S11494) for 30 min, and imaged using an Amersham ImageQuant 800 imager.

### Salt stability experiments

For the salt stability experiment in Fig. 1j, the nuclease footprinting assay was performed with the following modifications. MCM loading reactions (20 µl) were set up in assay buffer and were incubated (37°C, 30 min, 1,250 rpm). Then 10 µl of 3x sodium chloride in 1x assay buffer solutions were added to achieve the indicated sodium chloride concentrations, and the reaction was incubated (37°C, 15 min, 1,250 rpm). The reactions were diluted to a volume of 200 µl with solutions that adjusted all reaction buffers to 25 mM HEPES/KOH pH 7.6, 20 mM sodium acetate, 150 mM sodium chloride, 10 mM magnesium acetate, 1 mM DTT. Immediately, 10 µl of Benzonase nuclease (Sigma-Aldrich, E1014) was added and the mix incubated (37°C, 0.5 min, 1,250 rpm). Then 200 µl of the mix was transferred into 100 µl 3x Stop buffer and the mix incubated (37°C, 20 min, 1,250 rpm). The samples were extracted with an equal volume of phenol/chloroform/isoamyl alcohol, followed by two sequential rounds of ethanol precipitation and resuspension in TE. The salt stability experiments in Ext. Data Fig. 1d,e were performed in a similar way with the following modifications. MCM was loaded for 30 min in a single large reaction and split into two tubes before adding assay buffer or a 3x sodium chloride in 1x assay buffer solution to achieve the indicated salt concentration. Samples were taken at the indicated timepoints, diluted, treated with Benzonase, and transferred into Stop buffer. Dilution before Benzonase addition resulted in a salt concentration of 20 mM sodium acetate, 75 mM sodium chloride for the experiment in Ext. Data Fig. 1d, and in 20 mM sodium acetate, 112.5 mM sodium chloride for the experiment in Ext. Data Fig. 1e. For the experiment in Ext. Data Fig. 1e, instead of two sequential ethanol precipitations, the extracted aqueous phase was diluted 3-fold with TE and a single round of DNA precipitation with 50% isopropanol, 200 mM sodium chloride, 20 µg glycogen, followed by an 80% ethanol wash was performed.

### Geminin inhibition experiments

For the geminin inhibition experiment shown in Fig. 1i, geminin and CDT1 were mixed on ice prior to setting up the MCM loading reactions. Geminin/CDT1 mixes were prepared at 20x concentration in assay buffer + 200 mM sodium chloride. Reactions were then started by adding MCM (10x, assay buffer), ORC6 (20x, 200 mM NaCl), geminin-CDT1 mix (20x, 200 mM NaCl), and ORC1-5/CDC6 mix (40x, 650 mM NaCl) to a mix of DNA and ATP in assay buffer. For the ATPγS timecourse experiment in Ext. Data Fig. 7d geminin was added at minute 9 at 300 nM (2-fold excess over 150 nM CDT1ΔN).

### Nuclease footprinting assay with yeast proteins

Nuclease footprinting experiments with yeast proteins were performed at 30°C instead of 37°C. Reactions with yeast proteins contained 4 nM (27.5 ng/µl) plasmid DNA (10.6 kb, caesium chloride purified), 2 mM ATP, 100 nM yMcm2-7/Cdt1, 40 nM yCdc6, and 40 nM yOrc1-6. In the experiment shown in Fig. 3h human and yeast reactions were both carried out at 30°C.

### Experiments with human origin sequences

A 1,500 bp fragment of the origin at the human Lamin B2 locus, and a 2,398 bp fragment of the origin at the human Myc locus were cloned into the vector pBIG1c. The LamB2 origin fragment was amplified from human genomic DNA by PCR using oligonucleotides hOri-LamB_for (AACGCTCTATGGTCTAAAGATTTTACGTACATGCGGACCCCGCCTC) and hOri-LamB_rev (AACCCCGATTGAGATATAGATTTCAAGCCCTGCGTCCAGCGAG). The Myc origin fragment was amplified using oligonucleotides hOri-Myc_for (AACGCTCTATGGTCTAAAGATTTAAGCTTGTTTGGCCGTTTTAGGG) and hOri-Myc_rev (AACCCCGATTGAGATATAGATTTCTCGAGGCAGGAGGGGAG). The fragments were inserted into pBIG1c that was linearised with the restriction enzyme SwaI using Gibson assembly. For the experiment shown in Ext. Data Fig. 1f, a nuclease footprinting assay was performed using 27.5 ng/µl plasmid DNA of either pBIG1c, pBIG1c-LamB2_ori, or pBIG1c-Myc_ori.

### Bead-based recruitment assays

For the bead-based MCM recruitment assays in Ext. Data Fig. 1a,b double biotinylated DNA was generated by PCR with oligonucleotides Btn-1cOri_for (5’Biotin-GCTAACTGAAACACGGAAGGAGAC) and Btn-1cOri_rev (5’Biotin-TGGCTATGGCAGGGCTTG) on a human LaminB2 origin containing template (pBIG1c-LamB2_ori) resulting in a 1,823 bp PCR product. The DNA was coupled to magnetic Dynabeads M-280 Streptavidin (Invitrogen, 11206D) by incubation in buffer A (5 mM Tris pH 7.5, 0.5 mM EDTA, 1 M sodium chloride) for 30 min, 1.250 rpm, followed by 2 washes with buffer A, and 2 washes with buffer B (10 mM HEPES pH 7.6, 1 mM EDTA). Reactions were set up in bead-assay buffer (25 mM HEPES pH 7.6, 100 mM potassium glutamate, 10 mM magnesium acetate, 5% glycerol, 0.02% NP-40, 1 mM DTT) by mixing DNA beads with 5 mM ATP, 40 nM ORC1-5, 40 nM ORC6, 40 nM CDC6, 80 nM MCM2-7, and 80 nM CDT1.

Reactions were incubated for 30 min at 37°C with shaking at 1,250 rpm. Beads were then either washed with a low-stringency (LS) wash or washed more thoroughly with a sodium chloride containing buffer as indicated. A LS wash was performed by washing 2x for 20 s with shaking at 1,250 rpm using bead-assay buffer. A sodium chloride wash was performed by washing 2x for 2 min, 1,250 rpm with bead-assay buffer + sodium chloride as indicated, followed by a LS wash. For elution, DNA was degraded by nuclease treatment by resuspending the beads in bead-assay buffer + 2 mM calcium chloride, 600 u micrococcal nuclease (NEB, M0247S) and incubation at 37°C for 2 min with 1,250 rpm shaking. The eluate was analysed by SDS-PAGE using 3-8% Criterion XT Tris-Acetate gels (Bio-Rad) and XT Tricine running buffer (Bio-Rad), followed by silver staining using SilverQuest silver staining kit (Invitrogen, LC6070).

### Preparation of DNA templates for EM experiments

The DNA substrate used for EM imaging of wild type MCM loading was modified from the pGC209^14^ construct, containing two inverted ACS sequences spaced by 70 bp, by adding Widom601 and Widom603 strong positioning sequences at both ends. For the MCM AG mutant, Widom sequences were swapped for a suicide substrate for covalent *M.Hpa*II methyltransferase binding. The plasmid was synthesised by Eurofins and used for PCR amplification with the primer pairs NCP F / NCP R (for nucleosome reconstitution) or Gid70-MTRB F / Gid70-MTRB R (for methyltransferase capping).

Preparation of nucleosome- and *Hpa*II-flanked origins was carried out as described^28^. Briefly, amplified templates were purified by anion exchange chromatography on a 1 ml RESOURCE Q column (Cytiva), followed by ethanol precipitation. DNA pellets were resuspended in TE buffer, mixed with purified yeast histone octamers and subjected to dialysis with decreasing NaCl concentration to reconstitute nucleosomes^28,40^. The chromatinised construct was purified by size exclusion chromatography using a Superose 6 Increase 3.2/300 column (Cytiva). For the methyltransferase construct, DNA was incubated with *M.Hpa*II in 1:6 molar ratio at 30°C overnight in buffer 1 (50 mM potassium acetate, 25 mM Tris pH 7.5, 10 mM magnesium acetate, 1 mg/ml bovine serum albumin (BSA), 150 μM S-adenosyl-methionine (NEB)). The *M.Hpa*II–70bp–*M.Hpa*II construct was isolated by anion exchange chromatography using a 1 ml RESOURCE Q column (Cytiva).

NCP F - 5’ - (Des)-CGATAGAACTCGGGCCGCCCTGGAGAATCGCGGTGCCG – 3’

NCP R - 5’ - CCTGCACCCCAGGGACTTGAAGTAATAAGGAC – 3’

Gid70-MTRB F - atatatCC*GGcctgtATCTCGATTTTTTTATGTTTAGTTTCGC

Gid70-MTRB R - TGGGCGCC*GGAACTGGGTGCTGTaTTTTTATGTTTAGTTCG

(Des) – Desthiobiotin TEG

C* - 5-fluoro-2′-deoxycytosine

### Reconstitution of H. sapiens origin licensing for cryo-EM

For the hDH loading reaction, 45 nM of chromatinised DNA (nucleosome–Gid70– nucleosome) were incubated with 120 nM ORC1–5 1ΔN, 120 nM ORC6, 150 nM CDC6ΔN, 150 nM CDT1ΔN and 60 nM MCM2–7 in EM buffer (25 mM HEPES-KOH pH 7.6, 100 mM potassium glutamate,10 mM magnesium acetate, 1 mM DTT, 2 mM ATP) resulting in a final volume of 35 µl. Incubation was carried out for 30 minutes at 37°C and 1,250 rpm constant mixing.

The MCM recruitment reaction was established by substituting ATP with ATPγS and a chromatinised origin concentration of 70 nM.

### Sample preparation and data collection for negative stain EM

Dropout experiments shown in ED Fig. 2 were carried out with the same sample concentrations as described above, but using a final volume of 20 µl per reaction, and omitting one factor at a time. Reactions were diluted 4-fold in EM buffer where nucleotide was omitted. 300-mesh copper grids coated with a layer of continuous carbon (EM Resolutions, C300Cu100) were glow-discharged at 25 mA for 1 minute using a GloQube® Plus Glow Discharge System (Quorum), before applying 4 µl of the sample for 2 minutes. Grids were stained with two successive applications of 4 µl of 2% (w/v) uranyl acetate solution. Excess stain was blotted after 40 seconds using filter paper. Micrographs were collected using a FEI Tecnai G2 Spirit transmission electron microscope operated at 120 keV, equipped with a 2K x 2K GATAN UltraScan 1000 CCD camera. Data collection was carried out at a nominal magnification of 30,000x, yielding a pixel size of 3.45 Å at the specimen level, and a defocus range of -0.6 to -1.4 µm.

### Negative stain image processing

Negative stain images were processed using RELION-3.1^41^. Particles were picked using Topaz v0.2.5^42^. Contrast transfer function (CTF) parameters were estimated using Gctf v1.06^43^. Extracted particles were then subjected to reference-free 2D classification.

### Sample preparation and data collection for cryo-EM

UltrAuFoil R1.2/1.3 300-mesh grids (Quantifoil) were glow discharged at 40 mA for 5 minutes using a GloQube® Plus Glow Discharge System (Quorum), before applying 3 µl graphene oxide dispersion (10 ml graphene oxide flake dispersion (Sigma) diluted in 80 ml water; aggregates removed by centrifugation at 500 x g for 1 minute). Incubation was carried out for three minutes, followed by blotting of excess liquid and three successive washes with 20 µl droplets of water. After 1-2 hours drying at room temperature, 4 µl of the undiluted (ATP) reaction or the 3:1 diluted (ATPγS) reaction were applied to grids for 60 seconds at room temperature and 90% humidity in a Vitrobot Mark IV (Thermo Fisher). Grids were double-side plotted with force 0 for 5 seconds and immediately plunge frozen in liquid ethane. Micrographs were collected in counting mode using a pixel size of 1.08 Å on a Titan Krios transmission electron microscope with a K2 Summit direct electron detector and BioQuantum energy filter. A total electron dose of 49.28 e^-^/Å^2^ was used over 32 dose-fractioned movie frames and a total exposure time of 9.4 seconds. The defocus ranged from -1.0 to -2.5 µm. 3,589 movies were collected for the ATP reaction and a total of 31,569 movies for the ATPγS reaction.

### Cryo-EM image processing for the human DH loading reaction (ATP)

Image processing was performed using RELION-4.0b-GPU^46^ and cryoSPARC v3.3.2^44^ at different stages of the processing pipeline as indicated in ED Fig. 3. Beam-induced motion was accounted for by RELION’s implementation with 5 x 5 patches and CTF parameters were estimated using CTFFIND v4.1.10^45^. Particle picking was carried out using Topaz v0.2.4^42^, followed by particle extraction with a 440-pixel box and re-scaling to 110 pixel in cryoSPARC. 3,589 micrographs with 970,326 particles were selected based on the CTF fit resolution of 2.57–4.50 Å, CTF fit cross-correlation of 0.07–0.27 and median pick score of 20.18–43.55. Three rounds of reference-free 2D classification, *ab initio* reconstruction and heterogeneous refinement were conducted to identify 49,485 hDH particles. After particle extraction without downscaling and using a 400-pixel box, the particle stack was further cleaned up by 2D classification, which yielded 19,049 particles used to compute high-resolution hDH 2D class averages. *Ab initio* reconstruction, followed by homogeneous, non-uniform and local refinement with C2 symmetry resulted in map with 3.1 Å resolution. The particle stack was re-extracted, re-grouped, and cleaned from duplicate particles using RELION. 15,874 particles of the highest quality were isolated by 3D classification without alignment using a 320 Å mask and a regularisation parameter T of 4. Particles were polished^41^ subjected to 2D classification without alignment and 3D refinement imposing C2 symmetry. CTF parameters were optimised^41^ in three rounds (first, per-particle defocus, per-micrograph astigmatism; second, per-particle defocus, per-particle astigmatism, beamtilt; third, per-particle defocus, per-particle astigmatism, beamtilt, trefoil, 4th order aberrations) to yield a 3.3-Å resolution hDH structure. Homogeneous and non-uniform refinement with C2 symmetry in cryoSPARC resulted in the final map at 3.1 Å resolution.

To determine the structure of hSH from the same dataset, a new Topaz model was trained. 178,853 particles were extracted with a 440-pixel box, downscaled to 110 pixels. Smaller particles and contaminations were removed by 2D classification in cryoSPARC. Initial volumes were obtained by *ab initio* reconstruction with six classes. 37,396 hSH particles were then isolated in two rounds of heterogeneous refinement (ED Fig. 10) and re-extracted with a 400-pixel box. Homogeneous, non-uniform and local refinement with a mask encompassing the entire hSH yielded a 3.4 Å resolution structure. The same particles were re-extracted and 3D refined in RELION. CTF refinement (per-particle defocus, per-micrograph astigmatism), Bayesian polishing and one additional round of CTF refinement (per-particle defocus, per-particle astigmatism, beamtilt) were carried out. 25,069 high-resolution hSH particles were isolated by 3D classification. Homogeneous, non-uniform and local refinement with a mask around the hSH yielded 3.2 Å resolution in cryoSPARC.

### Cryo-EM image processing for H. sapiens MCM recruitment in the presence of ATPγS

Movies were corrected for beam-induced motion using RELION’s implementation with 5 x 5 patches in RELION-4.0b-GPU^46^ and CTF parameters were estimated using CTFFIND v4.1.13^45^. A Topaz model^42^ was trained and 1,334,277 particles were picked from 31,569 micrographs. Particles were extracted with a 416-pixel box, rescaled to 104 pixels. 629,241 MCM-containing particles (i. e. hOCCM, hSH, hMO*) were isolated using reference-free 2D classification. Initial volumes were generated using *ab initio* reconstruction in cryoSPARC. Low-pass filtered volumes were used for multi-reference 3D classification in RELION (ED Fig. 7). Particles contributing to the different complexes (114,995 hSH, 170,792 hOCCM and 203,088 hMO* particles) were re-extracted using a 400-pixel box without rescaling.

Homogeneous, non-uniform and local refinements with masks around the entire respective complex resulted in 3.6-Å resolution structure of the hSH, 4.0 Å hOCCM and 3.7 Å hMO* in cryoSPARC. Bayesian polishing was carried out for each particle stack individually in RELION^41^.

The hOCCM was then subjected to three rounds of CTF refinement (first, per-particle defocus, per-micrograph astigmatism; second, anisotropic magnification; third, per-particle defocus, per-particle astigmatism, beamtilt) followed by another round of Bayesian polishing in RELION. A mask around CDC6 and portions of ORC1–5 was generated to carry out focused 3D classification without alignment. This yielded 100,567 particles with good density for ORC–CDC6. 3D refinement in RELION was followed by homogeneous refinement in cryoSPARC. Local refinement with a mask around ORC–CDC6 was used to improve alignment on this part of the complex. 34,116 hOCCM particles with well resolved CDC6 density and 49,771 OC1M particles that lacked CDC6 were isolated by 3D classification without alignment in cryoSPARC. Both hOCCM and OC1M structures were locally refined to 3.8 and 4.1 Å, respectively, using a mask encompassing the entire complex.

Complexes containing single-loaded hexamers, hSH and hMO* particles were initially processed together (ED Fig. 7). 3D refinement in RELION using a mask around the MCM yielded a 3.8-Å hSH structure. Two rounds of CTF refinement (first, per-particle defocus, per-micrograph astigmatism; second, anisotropic magnification) and another round of Bayesian polishing was carried out. 135,742 hSH particles and 182,341 hMO* particles were subsequently separated by multi-reference 3D classification using the initial maps of hSH and hMO*, low-pass filtered to 30 Å. The hSH was refined in cryoSPARC (homogeneous, non-uniform, followed by local refinement) to 3.4 Å. Homogeneous and non-uniform refinement of the hMO* resulted in a consensus map solved to 3.6 Å resolution. The MCM portion of the complex was locally refined to 3.5 Å, while the ORC was refined to 4.0 Å.

### Preparation of MCM5AG DHs for cryo-EM

A 50-µl MCM double hexamer loading reaction was assembled using the protocol described in ‘Reconstitution of *H. sapiens* origin licensing for cryo-EM’. In short, 45 nM *M.Hpa*II– Gid70–*M.Hpa*II DNA capped at each end with TwinStrep-tagged *M.Hpa*II, was co-incubated with 90 nM ORC1–5 1ΔN, 150 nM CDC6ΔN, 150 nM CDT1ΔN and 60 nM MCM5AG/MCM2–7 in EM Buffer at 37°C under agitation. After 30 minutes, the loading reaction was diluted either by 40% or 75% with EM buffer and immediately used for cryo-EM.

UltrAuFoil R1.2/1.3 300-mesh grids were coated with a graphene oxide support as described above. 4 µL of diluted loading reaction were applied onto each grid in a Vitrobot Mark IV set to 22 °C and 90% humidity. After one minute on-grid incubation, grids were double-side blotted for 4.5 seconds with blot force 0 and plunged into liquid ethane. Grids were subsequently clipped and stored in liquid nitrogen until data collection.

### Cryo-EM data collection and image processing of the MCM5AG DH

13,203 movies were acquired from two grids at 92,000x magnification (1.58 Å/px) on a Glacios microscope equipped with a Falcon 3 direct electron detector operated in linear mode. A total dose of 50e^-^/Å^2^ (exposure time 1.12 seconds) and a defocus range of -1 to -2.5 µm (0.3 µm stepsize) were applied. Movie frames were aligned using RELION-4.0^46^ and CTF was estimated using GCTF v1.06^43^. 2,875,825 particles were picked using a pre-trained Topaz network^42^, extracted with a box size of 64 pixels (4x binned to 6.32 Å/px) and imported in cryoSPARC v4.0^44^ for 2 rounds of 2D classification. After 2D cleaning, two *ab initio* models were generated with C1 symmetry using a subset of 50,000 particles. All 379,664 particles were subjected to one round of heterogenous refinement against the two ab-initio reconstructions, followed by homogenous refinement in C1. Particles were re-extracted in RELION with a box size of 300 px (unbinned at 1.58 Å/px) and 2D-classified in cryoSPARC. The resulting 322,548 particles were used to generate an unbinned ab-initio reconstruction in C1, yielding a 5.76 Å map after non-uniform refinement in C1. 3D-classification in RELION without alignment using a regularisation parameter T of 200 distributed the particles into four classes with roughly equal populations, differing by presence of both, either or no Mcm6 WH. The best-resolved class (showing density for both Mcm6 WH domains) was non-uniform and locally refined in cryoSPARC with C2 symmetry, yielding a final resolution of 5.6 Å according to gold-standard FSC and the 0.143 criterion. Local resolution estimation was carried out in cryoSPARC.

### hDH model building

AlphaFold-Multimer^47^ was used to generate models of the ATPase tier (including the winged helix domains) of the hexameric human MCM2–7 assembly as well as the amino-terminal tier. Each model was rigid-body docked into one hexamer of the 3.1-Å resolution hDH map using UCSF ChimeraX v1.6.1^48^. Each chain was refined in Coot v0.9.8.1 EL^49^ and sections that could not be confidently built were deleted^48^. The models of the ATPase and amino-terminal tiers were then combined. ATP, ADP, magnesium and zinc ions were added in the pertinent sites. Idealised B-form DNA was first docked into the density of the double helix and then manually modified to account for the stretch of underwound and melted DNA within the N-terminal dimerisation interface. The model consisting of the MCM2–7 hexamer, ligands and DNA was adjusted using ISOLDE 1.6.0^50^, imposing ligand, secondary and base pairing restraints. This was followed by an iterative process of real space refinement with restraints on geometry, secondary structure, metal coordination and nucleic acid planarity in Phenix v1.21^51^ and manual adjustments in Coot and ISOLDE. To generate the hDH, a copy of the refined MCM hexamer was generated and rigid-body docked into the second hexamer of the hDH map. Clashes at the interface of the two hexamers were addressed using ISOLDE. The resulting model was subjected to real space refinement in Phenix.

### hSH model building

The atomic model of the hDH was rigid-body docked into the 3.2-Å cryo-EM map of the hSH using UCSF ChimeraX v1.6.1^52^. The second hexamer was deleted, nucleotides were inspected and the DNA was replaced by an idealised B-form duplex DNA using Coot v0.9.8.1 EL^53^. The model was refined with ISOLDE 1.6.0^50^ and Phenix real space refinement with restraints on geometry, secondary structure, metal coordination and nucleic acid planarity^51^.

### hMO* model building

To generate the model of the hMO* complex, the atomic structure of the hSH described above was docked into the map of the globally refined hMO* map using UCSF ChimeraX v1.6.1^48^. The atomic model of *H. sapiens* ORC6 was retrieved from the AlphaFold Protein Structure Database (accession code AF-Q9Y5N6-F1). To guide the positioning of ORC6, the *S. cerevisiae* MO complex (PDB entry 6RQC)^28^ was aligned with the human MCM in the map. Based on the alignment with the *S. cerevisiae* Orc6, the N-terminal cyclin box domain (residues 1–94) of human ORC6 was positioned next to the N-terminal domains of MCM2 and MCM6. The domain was then rigid-body docked into the density. The second cyclin box domain (residues 95–190) was docked into the adjacent density of the map that is positioned between MCM6 and ORC5. Docking solutions with the highest cross-correlation scores were chosen. The starting model of ORC1–5 was the open conformation of the human ORC1–5 complex (PDB code 7JPR)^54^. The model was docked into the locally refined map of ORC at 4.0 Å resolution. The DNA model of the *S. cerevisiae* MO complex (PDB entry 6RQC)^28^ was fit into the DNA density in the globally refined hMO* map. The DNA model was refined in Coot v0.9.8.1 EL^53^ applying all-molecule self-restraints 6.0 and DNA B-form restraints, before combining it with the DNA in the hSH. ORC6 was combined with the MCM and DNA into one PDB model. The two models of ORC6–MCM and ORC1–5 were iteratively refined with ISOLDE 1.6.0^50^ and Phenix real space refinement^51^ with restraints on geometry, secondary structure, metal coordination and nucleic acid planarity against the globally refined map of hMO* and the locally refined map of ORC, respectively. For illustration purposes, a composite map was generated using Phenix and the refined ORC6–MCM and ORC1–5 models were combined into one PDB model.

### hOCCM model building

To assemble the atomic model of hOCCM, the human ORC1–5–DNA model (PDB entry 7JPS), the ORC2 WH domain of the ORC1–5 model (PDB entry 7JPR)^54^ and the prediction of human CDC6 retrieved from AlphaFold Protein Structure Database (accession code AF-Q99741-F1) were rigid-body docked into the 3.8-Å cryo-EM map using UCSF ChimeraX v1.6.1^48^. Unstructured parts of the models, which were not visible in the hOCCM map, were deleted. The hSH model was docked into the map and each subunit was split and fitted independently into the density. The N-terminal tiers of MCM7, MCM3 and MCM5 were deleted from the atomic coordinates file as the local quality of region of the map was deemed insufficient for model building. The positioning of the ATPase tiers of the same MCM subunits was guided by the position in the hSH and no further adjustment of atomic positions was carried out due to the limited local resolution. The atomic models of the MCM3 and MCM7 WH domains were overlayed with equivalent domains in yOCCM. The atomic model of CDT1 was retrieved from the AlphaFold database (accession code AF-Q9H211-F1) and the three structured domains (residues 167–387, 418–440 and 441–546) were extracted from the model. Each domain was overlayed with yeast Cdt1 in yOCCM (PDB entry 5V8F)^29^. All models were combined into one PDB model and adjusted using Coot v0.9.8.1 EL^53^ and ISOLDE 1.6.0^50^. Models of ATPγS and magnesium were added in the ATPase active sites of MCM2–6, MCM6–4, ORC1–4, ORC4–5 and ORC5–3. A 39-mer idealised B-form duplex DNA was generated and fit into the density by applying all-molecule self-restraints 6.0 and DNA B-form restraints in Coot. The DNA coordinates were then merged with the hOCCM model before carrying out Phenix real space refinement with restraints on geometry, secondary structure, metal coordination and nucleic acid planarity^51^.

### Analysis of protein–DNA contacts

Protein–DNA contacts were analysed using the DNAproDB web-based visualisation tool^55,56^.

## Data availability

Data supporting the findings of this study are available within the paper and its Supplementary Information files. Cryo-EM density maps have been deposited in the Electron Microscopy Data Bank (EMDB) under the accession codes EMD-19618 (hDH), EMD-19625 (MCM5 AG DH), EMD-19619 (hSH), EMD-19620 (hMO*), EMD-19621 (locally refined ORC of hMO*), EMD-19622 (composite map of the globally refined hMO* and the locally refined ORC map), EMD-19623 (hOCCM), EMD-19624 (hOC1M). Atomic coordinates have been deposited in the Protein Data Bank (PDB) with the accession codes 8S09 (hDH), 8S0A (hSH), 8S0B (MCM–ORC6 of hMO*, which was modelled using the globally refined hMO* map), 8S0C (ORC1–5 of hMO*, which was modelled using the locally refined map of ORC), 8S0D (composite model of MCM–ORC6 and ORC1–5 docked into the composite map), 8S0E (hOCCM), 8S0F (hOC1M).

## Acknowledgements

We thank A. Alidoust, N. Patel, and D. Patel in the Structural Biology STP for yeast protein expression. We thank A. Nans, D. Benton, A. Purkiss and P. Walker from the Structural Biology STP for support with cryo-EM and computing. We are grateful to Marta Gross for the gift of yeast proteins. This work was supported by the Francis Crick Institute, which receives its core funding from Cancer Research UK (CC2002 and CC2009), the UK Medical Research Council (CC2002 and CC2009), and the Wellcome Trust (CC2002 and CC2009). This work was also funded by Wellcome Trust Senior Investigator Awards (219527/Z/19/Z) to JFXD, and European Research Council Advanced Grants (101020432-MeChroRep) to JFXD. AC receives funding from the European Research Council (ERC) under the European Union’s Horizon 2020 research and innovation programme (grant agreement no. 820102). FW has received funding from the European Union’s Horizon 2020 research and innovation programme under the Marie Sklodowska-Curie grant agreement No 844211. T.P. is the recipient of a Boehringer Ingelheim PhD fellowship. Work in T.C.R.M.’s lab is supported by a Novo Nordisk Fonden Hallas-Møller Emerging Investigator Grant (NNF22OC0073571), the Danish National Research Foundation (DNRF115) and the Carlsberg Foundation (CF21-0571). For the purpose of Open Access, the author has applied a CC BY public copyright licence to any Author Accepted Manuscript version arising from this submission.

## Author contributions

F.W., J.F.G., A.C. and J.F.X.D. conceived the study. F.W. cloned, expressed and purified human proteins and developed all biochemical assays. J.F.G., F.W. and T.C.R.M. performed negative stain work to develop and optimise the EM-based MCM loading assay. J.F.G. prepared cryo-EM grids, performed all cryo-EM data collection and image processing except for cryo-EM work on MCM5 AG, which was performed by T.P. J.F.G. performed all atomic model building and refinement. J.F.X.D. supervised the biochemical work and A.C. supervised the structural work. F.W., J.F.G., A.C. and J.F.X.D. wrote the manuscript with input from the other authors.

---

**Extended Data Fig. 1**

**a**, Bead-based MCM recruitment assay. A biotinylated PCR product was immobilised on Streptavidin-coated M-280 Dynabeads and used in full-length MCM loading reactions in the presence of ATP, ATPγS or no nucleotide. Beads were washed either with a low stringency wash (LS, 2x 20 sec 100 mM sodium acetate buffer) or washed with the same buffer containing additionally sodium chloride as indicated (2x 2 min, followed by a LS wash).

Beads were treated with micrococcal nuclease and the eluate analysed by SDS-PAGE and silver staining. **b**, As **a**, with ATP and omission of the indicated proteins. **c**, Representative negative-stain EM micrograph and 2D averages of a human MCM loading reaction on a short synthetic yeast origin flanked by nucleosomes. **d**, Salt stability of the DH footprint in a buffer containing 500 mM NaCl. MCM was loaded for 30 minutes and incubated with salt-containing buffers for the indicated time. Reactions were then diluted and treated with Benzonase. **e**, As **d**, with a buffer containing 750 mM NaCl. **f**, MCM loading on plasmid DNA with and without inserted origin DNA. Vec, pBIG1c. Vec-LamB2, pBIG1c containing a 1,500 bp fragment of the origin at the human Lamin B2 locus. Vec-Myc, pBIG1c containing a 2,398 bp fragment of the origin at the human Myc locus.

**Extended Data Fig. 2**

Representative negative-stain EM micrographs and 2D averages of human MCM loading reactions showing nucleotide and protein requirements for different reaction intermediates. **a**, Nucleosomes, ORC, MCM, OCCM, hMO* and DH classes were observed in the presence of ATP and all proteins. **b**, All complexes apart from DH were present when the slowly hydrolysable ATP analogue, ATPγS, was used. **c, d**, Without ORC6, DH classes were visible in the presence of ATP (**c**). No hMO* had formed when ORC6 was omitted, either in the presence of ATP (**c**) or ATPγS (**d**). **e,** DH formation depends on CDC6 in ATP conditions. **f**, In ATPγS conditions, OCCM-like and hMO* class averages were visible, even in the absence of CDC6. **g**, DH formation depends on CDT1 in ATP conditions. **h**, Similar to the experiments without CDC6, OCCM-like and hMO* class averages were detected when CDT1 was omitted in ATPγS conditions.

**Extended Data Fig. 3**

**a**, Representative cryo-electron micrograph of the human MCM loading reaction. **b**, 2D averages of nucleosomes, ORC, MCM, hMO* and DH were observed in the dataset. Box size 432 Å. **c**, Processing pipeline for the hDH. **d**, Angular distribution of the hDH structure. **e**, Fourier shell correlation plot for the hDH structure. **f**, Three rotated views and one cut-through view of the hDH structure, color-coded according to the local resolution as estimated in cryoSPARC.

**Extended Data Fig.4**

Protein–DNA interactions within the hSH, hMO*, hDH (asymmetric unit) and hOCCM.

**Extended Data Fig. 5**

**a,** Map of the AG mutant double hexamer coloured by subunit (see Figure for color code). **b,** Representative micrograph of a Mcm5 AG double hexamer loading reaction imaged at 92,000 x magnification. **c,** Processing pipeline of Mcm5 AG double hexamer sample. 13,203 acquired movies yielded a 5.56 Å structure following multiple rounds of 2D cleaning and 3D classification in RELION and cryoSPARC. **d,** Angular distribution of all particles contributing to the final reconstruction. **e,** Top (left) and side-views (centre) of the Mcm5AG double hexamer coloured by local resolution as calculated using cryoSPARC. A cut-through view (right) reveals untwisted DNA at the homo-dimerisation interface.

**Extended Data Fig. 6**

**a**, Nuclease footprinting assay with yeast proteins using ATP or ATPγS. **b**, Representative micrograph of the human MCM recruitment reaction using ATPγS. **c**, 2D averages of nucleosomes, ORC, MCM, OCCM and hMO* were observed in the dataset. The box size 449 Å. **d**, Angular distribution of the hOCCM structure. **e**, Fourier shell correlation plot for the hOCCM structure. **f**, Three rotated views and one cut-through view of the hOCCM structure, color-coded according to the local resolution as estimated in cryoSPARC. **g**, Angular distribution of the hOC1M structure. **h**, Fourier shell correlation plot for the hOC1M structure. **i**, Three rotated views and one cut-through view of the hOC1M structure, color-coded according to the local resolution as estimated in cryoSPARC.

**Extended Data Fig. 7**

Processing pipeline for the structures obtained in the ATPγS reaction.

**Extended Data Fig. 8**

**a,** Cryo-EM maps of the hOCCM, hOC1M and yOCCM. Cdt1 is colored in white to highlight the similar binding mode to MCM in the human and yeast complexes. **b**, Cut-through view of the hOCCM, hOC1M and yOCCM illustrating the interactions of the MCM WH domains with ORC–CDC6. MCM6 and MCM7 WH domains interact with ORC in all structures, while MCM3 is only visible when CDC6 is present. The MCM4 WH is visible only in yOCCM. **c**, Cartoon view of the WH interactions described in b. **d**, Nuclease footprinting assay showing reaction timecourses using ATPγS with (top) and without (bottom) addition of geminin at minute 9 after starting the reaction. **e**, Timecourse of Benzonase treatment (0.5 - 20 min) after a 30-minute reaction using ATPγS. A control was incubated for an additional 20 minutes followed by 0.5 min Benzonase treatment.

**Extended Data Fig. 9**

**a**, Angular distribution of the hSH structure that was obtained in the presence of ATP. **b**, Fourier shell correlation plot for the hSH structure. **c**, Three rotated views and one cut-through view of the hSH structure, color-coded according to the local resolution as estimated in cryoSPARC. **d**, Angular distribution of the globally refined hMO* structure. **e**, Fourier shell correlation plot for the globally refined hMO* structure. **f**, Three rotated views and one cut-through view of the globally refined hMO* structure, color-coded according to the local resolution as estimated in cryoSPARC. **g**, Locally refined ORC structure in the hMO*, color-coded according to the local resolution as estimated in cryoSPARC. **h**, Fourier shell correlation plot for the locally refined ORC structure. **i**, Locally refined MCM structure in the hMO*, color-coded according to the local resolution as estimated in cryoSPARC. **j**, Fourier shell correlation plot for the locally refined MCM structure.

**Exstended Data Fig. 10**

**a**, Processing pipeline for the hSH structure that was obtained from the human MCM loading reaction in the presence of ATP. **b**, Schematic representation of protein–protein interactions in hMO*, yMO, yMCM–Cdt1, hOCCM and yOCCM. In hMO*, the N-terminal cyclin box domain interacts with MCM2, MCM6 and ORC4, while the C-terminal cyclin box binds MCM6, MCM4 and ORC5. The resolution of the ORC3 specific insertion is limited. Thus, we cannot see the C-terminal alpha helix of ORC6 binding ORC3, which has been previously seen in the yeast and *Drosophila* ORC structures. The N-terminal cyclin box domain of yeast Orc6 binds similarly to Mcm2 and Mcm6, but it does not interact with Orc4. The C-terminal cyclin box domain of yeast Orc6 establishes contacts with Orc2 and Orc3. The C-terminal alpha helix of Orc6 also contacts Mcm5 and thereby bridges the Mcm2–5 DNA entry gate. In MCM–Cdt1, hOCCM and yOCCM, CDT1 establishes interactions with the same MCM subunits contacted by ORC6. In this context,the MCM2-5 gate appears accessible.

## References

1 Hu, Y. & Stillman, B. Origins of DNA replication in eukaryotes. Mol Cell 83, 352–372, doi:10.1016/j.molcel.2022.12.024 (2023).

2 Costa, A. & Diffley, J. F. X. The Initiation of Eukaryotic DNA Replication. Annu Rev Biochem 91, 107–131, doi:10.1146/annurev-biochem-072321-110228 (2022).

3 Bell, S. P. & Labib, K. Chromosome Duplication in Saccharomyces cerevisiae. Genetics 203, 1027–1067, doi:10.1534/genetics.115.186452 (2016).

4 Li, J. et al. The human pre-replication complex is an open complex. Cell 186, 98–111 e121, doi:10.1016/j.cell.2022.12.008 (2023).

5 Frigola, J., Remus, D., Mehanna, A. & Diffley, J. F. X. ATPase-dependent quality control of DNA replication origin licensing. Nature 495, 339–343, doi:10.1038/nature11920 (2013).

6 Fernandez-Cid, A. et al. An ORC/Cdc6/MCM2-7 complex is formed in a multistep reaction to serve as a platform for MCM double-hexamer assembly. Mol Cell 50, 577–588, doi:10.1016/j.molcel.2013.03.026 (2013).

7 Evrin, C. et al. A double-hexameric MCM2-7 complex is loaded onto origin DNA during licensing of eukaryotic DNA replication. Proc Natl Acad Sci U S A 106, 20240–20245, doi:0911500106 10.1073/pnas.0911500106 (2009).

8 Remus, D. et al. Concerted loading of Mcm2-7 double hexamers around DNA during DNA replication origin licensing. Cell 139, 719–730, doi:S0092-8674(09)01303-8 10.1016/j.cell.2009.10.015 (2009).

9. Lim, C., Miller, T., Tan, K.W., Talele, S., Early A., East P., Sanchez U., Dekker N.H., Costa, A., Diffley J.F.X. bioRxiv 10.1101/2024.01.10.575016 (2024).

10 Weissmann, F. et al. biGBac enables rapid gene assembly for the expression of large multisubunit protein complexes. Proc Natl Acad Sci U S A 113, E2564–2569, doi:10.1073/pnas.1604935113 (2016).

11 Vashee, S., Simancek, P., Challberg, M. D. & Kelly, T. J. Assembly of the human origin recognition complex. J Biol Chem 276, 26666–26673 (2001).

12 Siddiqui, K. & Stillman, B. ATP-dependent assembly of the human origin recognition complex. J Biol Chem 282, 32370–32383, doi:10.1074/jbc.M705905200 (2007).

13 Seki, T. & Diffley, J. F. X. Stepwise assembly of initiation proteins at budding yeast replication origins in vitro. Proc Natl Acad Sci U S A 97, 14115–14120, doi:10.1073/pnas.97.26.14115 (2000).

14 Coster, G. & Diffley, J. F. X. Bidirectional eukaryotic DNA replication is established by quasi-symmetrical helicase loading. Science 357, 314–318, doi:10.1126/science.aan0063 (2017).

15 Parker, M. W., Kao, J. A., Huang, A., Berger, J. M. & Botchan, M. R. Molecular determinants of phase separation for Drosophila DNA replication licensing factors. eLife 10, doi:10.7554/eLife.70535 (2021).

16 Parker, M. W. et al. A new class of disordered elements controls DNA replication through initiator self-assembly. eLife 8, doi:10.7554/eLife.48562 (2019).

17 Greiwe, J. F. et al. Structural mechanism for the selective phosphorylation of DNA-loaded MCM double hexamers by the Dbf4-dependent kinase. Nat Struct Mol Biol 29, 10–20, doi:10.1038/s41594-021-00698-z (2022).

18 McGarry, T. J. & Kirschner, M. W. Geminin, an inhibitor of DNA replication, is degraded during mitosis. Cell 93, 1043–1053 (1998).

19 Wohlschlegel, J. A. et al. Inhibition of eukaryotic DNA replication by geminin binding to Cdt1. Science 290, 2309–2312. (2000).

20 Tada, S., Li, A., Maiorano, D., Méchali, M. & Blow, J. J. Repression of origin assembly in metaphase depends on inhibition of RLF-B/Cdt1 by geminin. Nat Cell Biol 3, 107–113, doi:10.1038/35055000 (2001).

21 Frigola, J. et al. Cdt1 stabilizes an open MCM ring for helicase loading. Nature communications 8, 15720, doi:10.1038/ncomms15720 (2017).

22 Hu, Y. et al. Evolution of DNA replication origin specification and gene silencing mechanisms. Nature communications 11, 5175, doi:10.1038/s41467-020-18964-x (2020).

23 Lee, C. S. K. et al. Humanizing the yeast origin recognition complex. Nature communications 12, 33, doi:10.1038/s41467-020-20277-y (2021).

24 Vashee, S. et al. Sequence-independent DNA binding and replication initiation by the human origin recognition complex. Genes Dev 17, 1894–1908 (2003).

25 Li, N. et al. Structure of the eukaryotic MCM complex at 3.8 A. Nature, doi:10.1038/nature14685 (2015).

26 Evrin, C. et al. In the absence of ATPase activity, pre-RC formation is blocked prior to MCM2-7 hexamer dimerization. Nucleic Acids Res 41, 3162–3172, doi:10.1093/nar/gkt043 (2013).

27 Sun, J. et al. Cryo-EM structure of a helicase loading intermediate containing ORC-Cdc6-Cdt1-MCM2-7 bound to DNA. Nat Struct Mol Biol 20, 944–951, doi:10.1038/nsmb.2629 (2013).

28 Miller, T. C. R., Locke, J., Greiwe, J. F., Diffley, J. F. X. & Costa, A. Mechanism of head-to-head MCM double-hexamer formation revealed by cryo-EM. Nature 575, 704–710, doi:10.1038/s41586-019-1768-0 (2019).

29 Yuan, Z. et al. Structural basis of Mcm2-7 replicative helicase loading by ORC-Cdc6 and Cdt1. Nat Struct Mol Biol 24, 316–324, doi:10.1038/nsmb.3372 (2017).

30 Bleichert, F., Botchan, M. R. & Berger, J. M. Crystal structure of the eukaryotic origin recognition complex. Nature 519, 321–326, doi:10.1038/nature14239 (2015).

31 Chou, H. C. et al. The human origin recognition complex is essential for pre-RC assembly, mitosis, and maintenance of nuclear structure. eLife 10, doi:10.7554/eLife.61797 (2021).

32 Shibata, E. & Dutta, A. A human cancer cell line initiates DNA replication normally in the absence of ORC5 and ORC2 proteins. J Biol Chem 295, 16949–16959, doi:10.1074/jbc.RA120.015450 (2020).

33 Shibata, E. et al. Two subunits of human ORC are dispensable for DNA replication and proliferation. eLife 5, doi:10.7554/eLife.19084 (2016).

34 Bleichert, F. et al. A Meier-Gorlin syndrome mutation in a conserved C-terminal helix of Orc6 impedes origin recognition complex formation. eLife 2, e00882, doi:10.7554/eLife.00882 (2013).

35 Lin, Y. C. et al. Orc6 is a component of the replication fork and enables efficient mismatch repair. Proc Natl Acad Sci U S A 119, e2121406119, doi:10.1073/pnas.2121406119 (2022).

36 Lewis, J. S. et al. Mechanism of replication origin melting nucleated by CMG helicase assembly. Nature 606, 1007–1014, doi:10.1038/s41586-022-04829-4 (2022).

37 Bieniossek, C., Richmond, T. J. & Berger, I. MultiBac: multigene baculovirus-based eukaryotic protein complex production. Curr Protoc Protein Sci **Chapter** 5, Unit 5 20, doi:10.1002/0471140864.ps0520s51 (2008).

38 Vera Rodriguez, A., Frey, S. & Gorlich, D. Engineered SUMO/protease system identifies Pdr6 as a bidirectional nuclear transport receptor. J Cell Biol 218, 2006–2020, doi:10.1083/jcb.201812091 (2019).

39 Yeeles, J. T., Janska, A., Early, A. & Diffley, J. F. X. How the Eukaryotic Replisome Achieves Rapid and Efficient DNA Replication. Mol Cell 65, 105–116, doi:10.1016/j.molcel.2016.11.017 (2017).

40 Luger, K., Rechsteiner, T. J. & Richmond, T. J. Preparation of nucleosome core particle from recombinant histones. Methods Enzymol 304, 3–19 (1999).

41. Zivanov, J., Nakane, T. & Scheres, S. H. W. Estimation of high-order aberrations and anisotropic magnification from cryo-EM data sets in RELION-3.1. IUCrJ 7, 253–267, doi:10.1107/S2052252520000081 (2020).

42 Bepler, T. et al. Positive-unlabeled convolutional neural networks for particle picking in cryo-electron micrographs. Nat Methods 16, 1153–1160, doi:10.1038/s41592-019-0575-8 (2019).

43 Zhang, K. Gctf: Real-time CTF determination and correction. Journal of structural biology 193, 1–12, doi:10.1016/j.jsb.2015.11.003 (2016).

44 Punjani, A., Rubinstein, J. L., Fleet, D. J. & Brubaker, M. A. cryoSPARC: algorithms for rapid unsupervised cryo-EM structure determination. Nat Methods 14, 290–296, doi:10.1038/nmeth.4169 (2017).

45 Rohou, A. & Grigorieff, N. CTFFIND4: Fast and accurate defocus estimation from electron micrographs. Journal of structural biology 192, 216–221, doi:10.1016/j.jsb.2015.08.008 (2015).

46 Kimanius, D., Dong, L., Sharov, G., Nakane, T. & Scheres, S. H. W. New tools for automated cryo-EM single-particle analysis in RELION-4.0. Biochem J 478, 4169–4185, doi:10.1042/BCJ20210708 (2021).

47 Jumper, J. & Hassabis, D. Protein structure predictions to atomic accuracy with AlphaFold. Nat Methods 19, 11–12, doi:10.1038/s41592-021-01362-6 (2022).

48 Pettersen, E. F. et al. UCSF ChimeraX: Structure visualization for researchers, educators, and developers. Protein science : a publication of the Protein Society 30, 70–82, doi:10.1002/pro.3943 (2021).

49 Caspari, T. & Carr, A. M. DNA structure checkpoint pathways in Schizosaccharomyces pombe. Biochimie 81, 173–181. (1999).

50 Croll, T. I. ISOLDE: a physically realistic environment for model building into low-resolution electron-density maps. Acta Crystallogr D Struct Biol 74, 519–530, doi:10.1107/S2059798318002425 (2018).

51 Liebschner, D. et al. Macromolecular structure determination using X-rays, neutrons and electrons: recent developments in Phenix. Acta Crystallogr D Struct Biol 75, 861–877, doi:10.1107/S2059798319011471 (2019).

52 Pettersen, E. F. et al. UCSF Chimera--a visualization system for exploratory research and analysis. Journal of computational chemistry 25, 1605–1612, doi:10.1002/jcc.20084 (2004).

53 Casanal, A., Lohkamp, B. & Emsley, P. Current developments in Coot for macromolecular model building of Electron Cryo-microscopy and Crystallographic Data. Protein science : a publication of the Protein Society 29, 1069–1078, doi:10.1002/pro.3791 (2020).

54 Jaremko, M. J., On, K. F., Thomas, D. R., Stillman, B. & Joshua-Tor, L. The dynamic nature of the human Origin Recognition Complex revealed through five cryoEM structures. eLife 9, doi:10.7554/eLife.58622 (2020).

55 Sagendorf, J. M., Berman, H. M. & Rohs, R. DNAproDB: an interactive tool for structural analysis of DNA-protein complexes. Nucleic Acids Res 45, W89–W97, doi:10.1093/nar/gkx272 (2017).

56 Sagendorf, J. M., Markarian, N., Berman, H. M. & Rohs, R. DNAproDB: an expanded database and web-based tool for structural analysis of DNA-protein complexes. Nucleic Acids Res 48, D277–D287, doi:10.1093/nar/gkz889 (2020).

